# From Skin to Cortex: End-to-End Spiking Neural Network Simulation of Tactile Information Flow

**DOI:** 10.64898/2025.12.03.692094

**Authors:** Nicolas Rault, Tea Tompos, Emma F. Marx, Fleur Zeldenrust, Levent Degertekin, Tansu Celikel

**Affiliations:** Donders Institute for Brain, Cognition, and Behaviour, Radboud University, Nijmegen, the Netherlands; School of Psychology, Georgia Institute of Technology, Atlanta - GA, USA; Department of Mechanical Engineering, Georgia Institute of Technology, Atlanta - GA, USA; Georgia Tech-CNRS IRL 2958, Georgia Institute of Technology - Europe, Metz, France

**Keywords:** Spiking neural network, information processing, sensory integration, neuromorphic computing, tactile navigation

## Abstract

Autonomous systems and neuroprosthetic devices demand real-time tactile processing under strict energy and latency constraints. Designing these systems using neuromorphic principles, where communication is event-based and node activity is sparse, could improve their speed and energy efficiency. Here we present an end-to-end spiking neural network model of the ascending tactile pathway, from mechanoreceptors in the skin in humans (or whisker follicles in rodents) to cortical neurons, that operates in an event-driven neuromorphic fashion. The model comprises distinct anatomical stages, (1) three types of mechanoreceptor afferents, (2) trigeminal ganglion, (3) brainstem, (4) thalamic nucleus, and (5) three cortical layers, connected in a feed-forward hierarchy. We demonstrate the model’s responses to both rodent whisker deflection and human fingertip skin displacement, using information-theoretic analysis, pairwise correlation, and stimulus decoding at each layer. Our results show that tactile information is efficiently encoded and transformed at each stage: stimulus features are represented with high fidelity and reduced redundancy as signals ascend. Notably, simple linear or Bayesian decoders can reliably classify stimulus features from single-neuron activity in the thalamus and cortex for low-noise inputs, highlighting the emergence of robust neural representations. This open-source model is the first to include the mechanosensory periphery in a full tactile pathway simulation, enabling researchers to study how perturbations at any stage affect tactile encoding. Moreover, the network is well-suited for deployment on low-power, real-time neuromorphic hardware, facilitating the development of multi-layer signal processing and tactile navigation algorithms.

## Introduction

Autonomous and human-in-the-loop systems alike require real-time processing of sensory inputs under strict energy and latency constraints. This is particularly challenging for tactile sensing, where touch information contains rich and rapidly changing spatio-temporal features [1, 2]. Current tactile systems often rely on high-frequency sampling with constant rates [3], regardless of the rate dynamics of external events, resulting in inefficient sensory representations. Moreover, these systems use centralized computation, suffering from poor scalability for assistive portable devices or embedded platforms. For instance, both prosthetic devices and teleoperated machinery suffer from these limitations, as they require real-time integration of asynchronous events with high-resolution [4]. In contrast, biological tactile systems, such as finger-based touch in primates or whisker-based touch in rodents, operate asynchronously and integrate information over parallel streams [2], offering a compelling alternative to systems designed using traditional principles. To advance state-of-the-art tactile systems, here, we employ a neuromorphic approach to develop a touch-encoding spiking neural network (SNN) model based on the anatomical organisation of the ascending somatosensory pathway. We model mechanosensitive cells as an input layer, which converts distinct analog-based stimulus features into digitized information (neural spikes), generating sparse and event-driven patterns [5, 6]. The spike-based signal is then relayed to the downstream network layers, communicating relevant stimulus features while minimizing redundancy [7].

Parts of the ascending somatosensory pathway of rodents [8] have been previously modelled using SNNs. For instance, Furuta et al. (2021) [6] modelled the peripheral pathway stages by representing different mechanosensitive cells within a whisker follicle. Huang et al. (2022, 2025) [10, 11], Sharp et al. (2014) [12] and Tompos et al. (2025) [23] modelled the central pathway stages, specifically representing multiple somatosensory cortical layers, using various levels of biological realism. These biologically grounded, i.e. neuromorphic, models were used to address how the sensory system encodes tactile stimuli and transfers information across layers, advancing our understanding of the tactile system’s function, completing and building on the experimental observations (ref).

To better understand the tactile system and enhance current sensory information processing devices, we must address two significant gaps. First, the peripheral and central stages of the somatosensory system remain unintegrated, as separate models across different platforms and scales. Second, as mentioned earlier, the SNNs are a great alternative to traditional tactile systems; however, there exists no common pipeline to evaluate the capacity of different neuromorphic architectures to integrate sensory information, their encoding strategies, or how network dynamics affect tactile representations across layers. This prevents the comparison of neuromorphic tactile architectures and slows the development of deployable solutions.

To address both of these gaps, we developed a multi-layer SNN model of the full ascending tactile pathway, along with an analysis pipeline for quantifying representational efficiency across layers, suitable for any neuromorphic, spike-based system. In summary, the key components of our effort are

1. Mechanoreceptor Layer: We modelled the dynamics of three types of mechanoreceptors in the model’s input layer, which capture all key dimensions of a tactile stimulus (e.g. amplitude, frequency, direction).
2. Multi-Layer Network: The spikes generated in the input layer propagate through a biologically realistic, multi-layer SNN architecture. Each layer captured the anatomical organization and the neuronal dynamics of one stage along the ascending pathway, spanning the trigeminal ganglion (TG), brainstem (BS), thalamus (VPM), and cortical layer (L) 4, L3 and L2.
3. Information Analytical Tools: We utilized information theoretic methods to measure spike-based information transfer and redundancy (mutual information and neural correlation analyses), which we applied to the spiking activity from every network layer.
4. Decoding Evaluation: We developed three distinct decoders to assess the quality and robustness of tactile representations at each layer by retrieving stimulus features from neural activity within layers.

This end-to-end, skin-to-cortex SNN model and accompanying analyses tools are fully open-source, allowing for straightforward modification of neuron models, connectivity patterns, and parameters. The model provides a foundation for exploring how network structure, cell-type properties, and synaptic dynamics shape tactile encoding. Furthermore, our work is readily deployable to neuromorphic hardware, with potential for low-power, real-time tactile processing applications.

## Methods

We describe here how the model was derived and how the resulting neural spiking activity was analyzed. We constructed a model of the ascending somatosensory pathway (Figure 1) beginning with three populations of mechanoreceptors and reflecting the known biological architecture. The input layer comprised SA1, SA2, and RA receptor populations, fitted to electrophysiological recordings (see section 2.3). The network was driven by two classes of stimuli. First, whisker deflection was simulated using a biomechanical model of the whisker fitted to recorded whisker kinematics (section 2.1.1), from which forces and torques at the whisker base were extracted to replicate the input received by mechanosensitive cells in the follicle [6]. Second, human tactile input was obtained from skin displacement at the fingertip measured during focused ultrasound stimulation (section 2.1.2). The spiking activity in the mechanoreceptors as a result of the stimuli propagates through successive layers representing the trigeminal ganglion (TG), brainstem (BS), thalamus (VPM), and the primary somatosensory cortex (bfd: barrel field). Information transfer across the pathway was quantified through information-theoretic metrics (section 2.10), and the coding structure was characterized by calculating the correlations between neural populations (section 2.11). Finally, we assessed the quality of stimulus representation by training decoders to recover tactile parameters from population and single-neuron activity (section 2.12).

**Figure 1.**
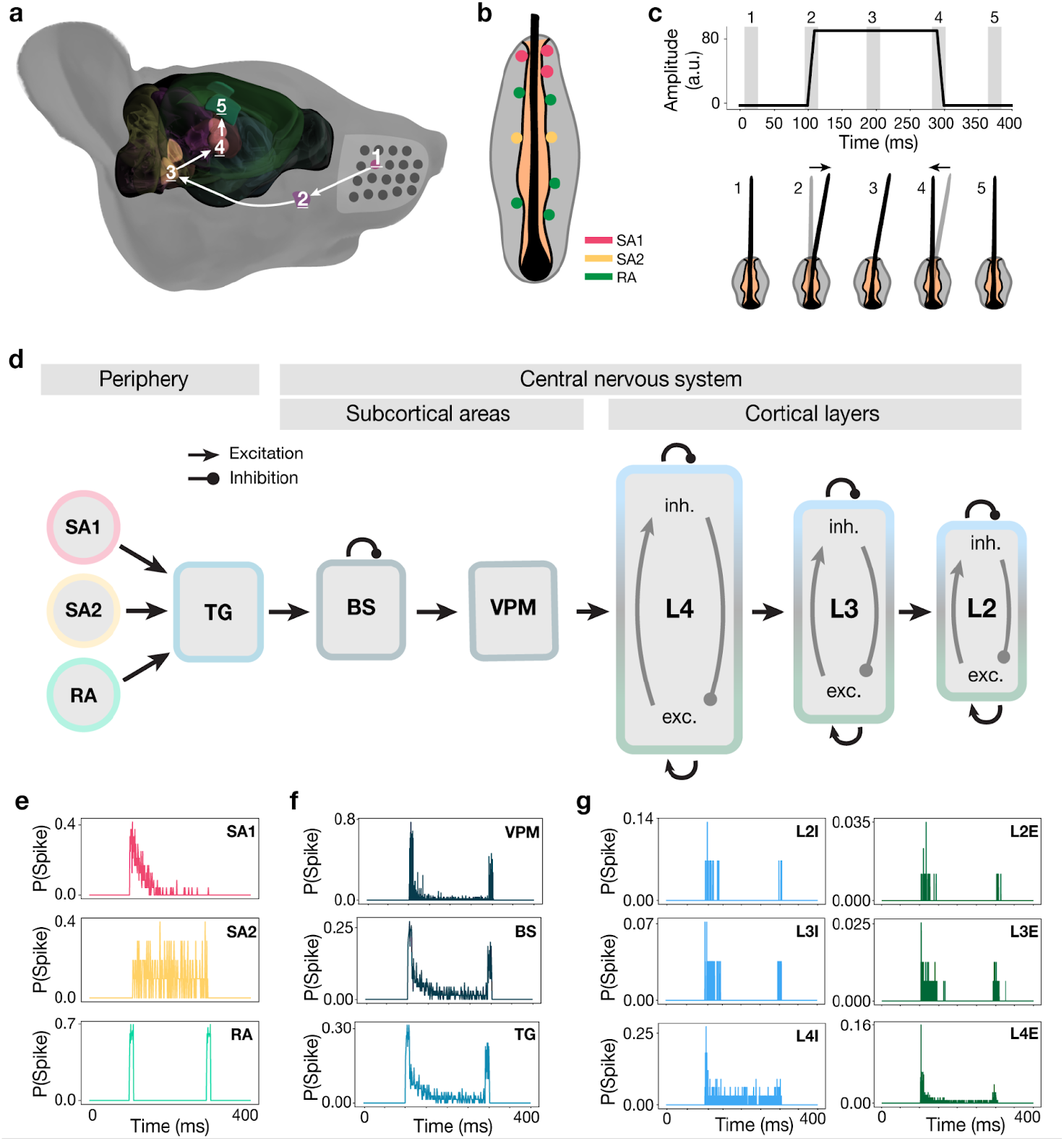
Neural responses from mechanoreceptors in the periphery to the superficial cortical layers in the central nervous system triggered by the ramp-and-hold stimulus. **(a)** Side view of a mouse head highlighting the pathway starting in the whisker follicle (1), going to the trigeminal ganglion (2), the brainstem (3), the thalamus (4), and finally the cortex (5). **(b)** Side-view of a whisker follicle ((1) in a) with three types of mechanoreceptors: slowly adapting type I (SA1, red) and II (SA2, yellow), and rapidly adapting (RA, green). **(c)** Time-varying amplitude (a.u.) of the ramp-and-hold stimulus composed of multiple phases (grey shades): (1) no stimulus, (2) ramp-on, (3) plateau, (4) ramp-off, and (5) no stimulus. The horizontal axis represents time in ms, and the vertical axis represents the amplitude of whisker deflection. Whisker position during each phase (1-5) is illustrated below (Figure 2). **(d)** The architecture of the spiking neural network model of the ascending somatosensory pathway. The model contains three excitatory populations representing mechanoreceptors (see panel **b**): SA1 (24 neurons), SA2 (12 neurons) and RA (36 neurons), the trigeminal ganglion (TG; 70 neurons), brainstem (BS; 70 neurons), ventral posteromedial (VPM; 70 neurons) thalamic nucleus with excitatory neurons, and three cortical layers: L4 (188 excitatory, 33 inhibitory), L3 (158 excitatory, 28 inhibitory) and L2 (86 excitatory, 15 inhibitory). Cortical layers contain excitatory (exc.) and inhibitory (inh.) populations. Lines with arrowheads represent excitatory connections. Lines with circleheads represent inhibitory connections. **(e)** The normalized (number of spikes divided by the population size) number of spikes for a population of SA1 (top), SA2 (middle) and RA (bottom) mechanoreceptors as a response to the ramp-and-hold stimulus (shown in c). The horizontal axis represents the time, and the vertical axis represents the normalized number of spikes. **(f)** Same as in e, but for VPM (top), BS (middle), and TG (bottom) populations. **(g)** Same as in e, but for inhibitory (left column) and excitatory (right column) L2 (top), L3 (middle) and L4 (bottom) populations.

All model code and analysis scripts are available on GitHub at: https://github.com/Nrault/StimulusRepInAscendingPathway.

### 2.1 Stimuli

#### 2.1.1 Simulated whiskers

The first component of the whisker-to-cortex model is the whisker itself. To accurately grasp the behavior of a whisker, we rely on the WhiskIt model [16]. In this model, each whisker is composed of 20 links held together by springs for which the elasticity and dampening coefficients were optimized to match the kinematics of recorded whisker trajectories during whisking behavior. For the current simulation, we focus on a single whisker in the ramp-and-hold paradigm. The ramp-and-hold paradigm is a passive perception paradigm where a pole is pushed against a single whisker during a ramp-on phase, held into position during the hold phase, and released back to its initial position during a ramp-off phase (Figure 1.c and Figure 2). This paradigm allows the manipulation of the direction with which the pole is pushed against the whisker (direction), its distance along the whisker axis to the base (distance), the speed at which the pole moves (speed), and how much pressure the pole applies to the whisker (amplitude), as shown in Figure 2. We also implemented a paradigm (frequency paradigm) where the pole is held against the tip of the whisker and vibrates at different frequencies. Thus, in this set of experiments, we manipulated direction, distance to the whisker basis, slope speed of the ramp-and-hold stimulus, amplitude of whisker deflection, and frequency of vibration. For the ramp-and-hold stimuli, each whisker stimulation lasted for 250ms, of which the hold phase lasted 100ms. For each set of ramp-and-hold stimuli (but not the frequency paradigm), the same set of initial parameters was used: for the first stimulus, the pole was positioned at 10 mm from the targeted whisker and moved at a speed of 0.3 m/s, with an angle of 0° around the x-axis (along the whisker). The amplitude of the stimulus was 1 mm, and it was held against the whisker for 100 ms. For the frequency paradigm the pole was placed against the tip of the whisker for the entire trial duration.

**Figure 2.**
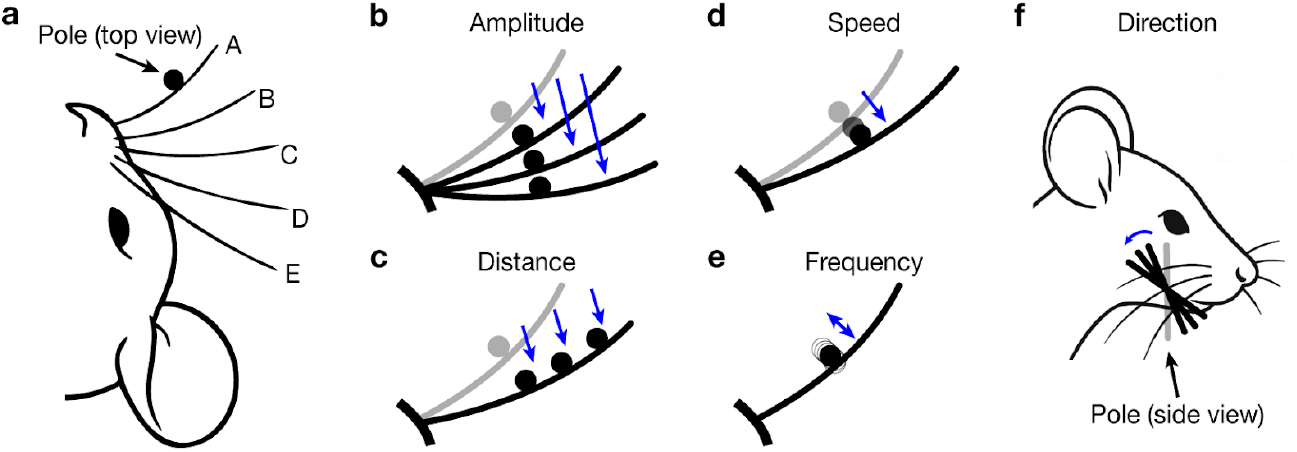
Stimulus paradigms for simulated whisker deflection experiments. **(a)** Baseline setup for whisker deflection: A pole is pushed toward the whisker A of the mouse, held in position, and moved back (ramp-and-hold). **(b)** Amplitude variation: (one whisker close-up): from the baseline (in grey), the amplitude of the deflection was varied. **(c)** Distance variation (one whisker close-up): from the baseline (in grey), the distance between the pole and the whisker base was varied. **(d)** Speed variation (one whisker close-up): from the baseline (in grey), the speed of movement of the pole toward the whisker during the ramp phases was varied. **(e)** Frequency variation (one whisker close-up): In the frequency paradigm, the pole was held against the whisker and vibrated at a constant frequency for one trial. Across trials, this frequency was changed. **(f)** Direction variation: from the baseline (in grey), we varied the direction with which the pole was pushed against the whisker.

For sets of stimuli where the maximum amplitude was varied, the maximum amplitude was varied from 1 mm to 20 mm with a step of 1 mm. For sets of stimuli where the direction was varied, the pole was oriented from 0° to 350° with steps of 10°). For sets of stimuli where the distance to the whisker basis was varied, we targeted each of the 20 links that compose the whisker once (see –ref Whiskit paragraph and Figure 2). For sets of stimuli where the slope speed was varied, the pole moved from 0.25 to 0.35 m/s with steps of 0.01 m/s. Finally, for sets of stimuli where the frequency was varied, the pole was placed against the last link of the whisker and vibrated with a frequency of 2 to 32 Hz with steps of 2 Hz).

From the simulations described above, we extracted the amplitude of the deflection of the base of the whisker as follows:

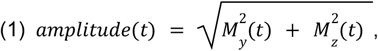

(equation from Yang et al., (2016) [17]).where *M*_*y*_ denotes the bending moment at the whisker basis on the y-axis (see below), and *M*_*z*_ denotes the bending moment at the whisker basis on the z-axis (see below). We define the y-axis as oriented toward the top of the head of the mouse, the z-axis toward its snout, and the x-axis, perpendicular to the z and y-axes toward the whisker tip at rest. Here, we assume that the moment along the x-axis has a negligible effect on the base deflection, and we therefore only consider the YZ moments. A moving average with a window size of 11ms, similar to [16], was applied to the resulting amplitude with original timestep of 1ms to reduce whisker oscillations.

#### 2.1.2 Recorded fingertips

Fingertip skin displacement during ultrasonic stimulation was recorded using a Polytec PDV-100 laser Doppler vibrometer (Polytec GmbH, Waldbronn, Germany). The ultrasound transducer (mounted in a chamber with a transparent bottom) had a central aperture allowing optical access to the fingerpad. The vibrometer was positioned approximately 40 cm below the fingertip and focused its laser through the chamber window and transducer aperture onto the skin. It measured the vertical surface motion of the fingerpad in real time. The vibrometer’s differential output (measured relative to a reference channel) was acquired at a 125 kHz sampling rate, providing a high-resolution displacement signal for each stimulus trial.

The recorded displacement signals were preprocessed. First, any baseline offset was subtracted from the raw displacement waveform. The 125 kHz signals were then subsampled at 1 kHz to match the model’s time step. To mimic the fingertip’s viscoelastic damping, an 11 ms moving-average filter was applied, corresponding to the skin’s fast viscoelastic response on the order of ~10 ms [18]. Next, the smoothed displacement trace was full-wave rectified (taking the absolute value) to capture the magnitude of skin motion independent of direction. Finally, the rectified displacement profiles were normalized across trials and scaled appropriately. The result was used as *amplitude*(*t*) input into the spiking neural network model. The full procedure is illustrated in the supplemental Figure S11.

Skin displacement data were recorded with 160 different stimulation amplitudes for a pulse with a fixed duration.

### 2.2 Input Layer

The amplitudes that were extracted from either the whisker deflections or the skin displacement recordings were directly applied to the mechanoreceptors. This input layer is composed of three mechanoreceptor populations representing Slowly Adapting type 1 (SA1), Slowly Adapting type 2 (SA2), and Rapidly Adapting (RA) mechanoreceptors as found in the literature [19]. The SA1 cells respond with a surge of activity followed by a decrease in their activity slightly after the onset of a stimulus, toward a plateau phase activity with an irregular spiking pattern [5]. SA2 cells have a slower decay time than SA1 cells and a higher activity level during the plateau phase [20]. RA cells respond to the transient part of the stimulus (ramp on and off) [20]. These cells’ recordings mainly differ in their adaptation rate [19]. To model these input cells, we rely on the Adaptive Exponential Integrate and Fire (AdEx) neuron model [21], which explicitly implements an adaptive current. The basis of the model is described in the equations below (2, 3, and 4):

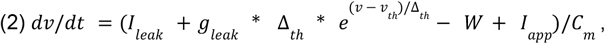

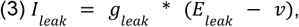

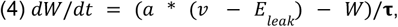

Where *I*_*leak*_ represents the leak current,*g*_*leak*_ the leak conductance, Δ_*th*_ the action potential slope factor, *v* the membrane potential, *v*_*th*_ the action potential threshold, *W* the adaptive current, *I*_*app*_ the applied current, *C*_*m*_ the capacitance, *E*_*leak*_ the leak reverse potential, *a* the strength of adaptation and τ the adaptation time constant. The values of each parameter for each receptor population are described in Supplemental Table S1.

Upon application of a current *I*_*app*_ to the cell, the membrane potential *v* will increase; if it reaches a spike threshold *v* ^*th*^ (−30mV), it is reset to its resting potential *v*_*r*_ (−45mV), and the adaptive current *W* is increased by a value *b*. If the threshold is not reached, then the membrane potential and the adaptive current both follow an exponential decay toward their resting values. In the case of several spikes, the adaptive current increases logarithmically before reaching an asymptote where the adaptation saturates (4). We used this model for our receptors and modified the applied current for each cell type according to the descriptions in the following sections.

To ensure that the receptive fields of the mechanoreceptors cover the entire space surrounding a whisker (i.e., a deflection of the whisker in any direction results in activation of a mechanoreceptor), the receptive fields were modeled as follows: each mechanoreceptor *i* has a cosine-shaped receptive field, where the center occupies a location *p*_*i*_ distributed in a circle on the yz plane around the whisker.

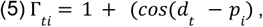

where t ∈ T denotes the duration of the simulation, *d*_*t*_ the direction of the whisker deflection at time t, *p*_*i*_ the position of mechanoreceptor i, and Γ_*ti*_ ∈ [0; 1] the mechanoreceptor *i* selectivity as a function of direction at time *d*_*t*_. The parameter *p*_*i*_ that gives each receptor *i*’s maximum Γ_*ti*_ was unique for each receptor and was drawn from a Gaussian distribution centered around 0 degrees (toward the snout of the mouse) and 180 degrees (toward the tail of the mouse) with a standard deviation of 90 degrees. To model the receptive field of a mechanoreceptor with respect to fingertip stimulation, we used a 2D Gaussian function, centered around *x*_*i*_ and *y*_*i*_,

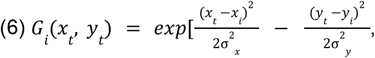

with σ_*x*_ and σ_*y*_ the standard deviation on the x-axis (width of the fingertip) and y-axis (length of the fingertip), respectively. *x*_*t*_ and *y*_*t*_ represent the position of the stimulus on the fingertip on the x and y axes at a given time *t. x*_*i*_ and *y*_*i*_ represent the position of the mechanoreceptor *i* on the fingertip along the x and y axes. With the whisker (Γ_*ti*_) and fingertip (*G*_*i*_ (*x*_*t*_, *y*_*t*_)) receptive fields defined, we can now describe the neural input (*I*_*app*_*(t)*, equation (2)) over time to each mechanoreceptor for each whisker displacement time series or to each fingertip stimulation (*amplitude(t)*).

The response of a SA1 receptoris mostly modulated by the direction of the whisker deflection, and not so much by the amplitude of the displacement [2, 6, 20, 22]. Therefore, for SA1 receptors, we assume an ON/OFF relationship to differences in amplitude of whisker deflection; that is, the stimulus current *I*_*app*_ is multiplied by 1 if there is a stimulus and 0 otherwise (equations 7, 8). The activation threshold *H*(*t*) to estimate whether a stimulus was present (sensitivity) was hand-tuned and set to 0.01 (amplitude, a. u.) to respond to the onset of the stimulus while remaining silent for noise induced amplitude:

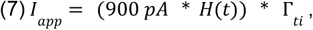

with t ∈ T, the time point in the simulation, and *i* the ID of the mechanoreceptor. The activation of the response due to ampltidue is modelled by a Heaviside step function:

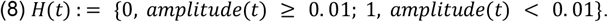

For the fingertip stimuli, SA1’s equations were adapted as follows:

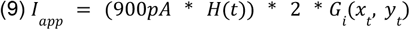

Since SA2 receptors are sensitive to both the direction and amplitudeof whisker deflection [2, 6, 20, 22], we, assumed a logarithmic increase of theresponse as a function of the whisker deflection amplitude:

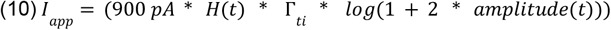

For fingertips, we adapted this equation to

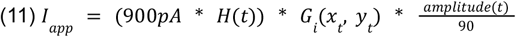

We assume that RA receptors are sensitive to a change in amplitude [2, 6, 20, 22].

Therefore, we modelled their activation as the derivative in time of the amplitude:

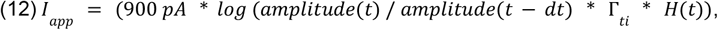

For fingertip stimulation, the RA mechanoreceptor equations were adapted to s:

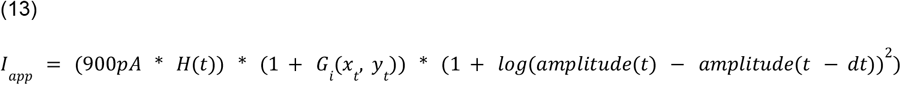

### 2.3 Fitting of the mechanoreceptor model parameters

The parameters (*a, b*, and τ) used in the receptor models SA1, SA2, and RA were fitted to recordings performed by Sonekatsu and Gu [19] in 2019 (supplemental Figure S16). The authors extracted a whisker follicle from a mouse and recorded the response of the receptor to a ramp-and-hold stimulus directly. From there, they computed the instantaneous firing rate (1/ISI, interspike interval). To fit the AdEx model parameters to the recordings, we used the CMAES algorithm from the CMA python toolbox, a covariance-matrix-based evolutionary algorithm that relies on the covariance between the parameters of the model and the error function (here a mean squared error, MSE between the instantaneous firing rate of the modeled receptors and the instantaneous firing rate of Sonekatsu and Gu’s recordings [19]). With this, we optimized the adaptive parameters of the model (i.e., *a, b*, and *τ*, see Supplemental Table S1), allowing us to replicate the adaptation rate of the different populations of receptors. The parameters of the RA receptors were hand-fitted, as the number of data points was too low to allow the model to train. We thus changed the value of the adaptation current (*a, b*, and *τ*) so that RA responds to the transient part of the stimulus.

### 2.4 Network models for the early stages of the central nervous system

Like the mechanoreceptors in the input layer, the trigeminal ganglion (TG) and brainstem (BS) neurons were also modeled using adaptive exponential integrate and fire models (Adex with parameters *a* and *b* set to 0 to inactivate the adaptation). TG neurons receive one-to-one connections (see Supplemental Table S2) from the receptor layer to aggregate neurons from the input layer while keeping the identity of the cells (SA1, SA2, and RA), with each presynaptic action potential resulting in an instantaneous increase in the postsynaptic membrane potential that was set so that a TG neuron responds to the activity of a single receptor (Supplemental Table S3). BS neurons receive a probabilistic connection pattern. Each BS neuron has a probability of 0.01 to receive an input from a TG neuron. Similarly, each BS neuron has a probability of 0.01 to send an inhibitory connection to another BS neuron (Supplemental Table S3). Excitatory synapses are modeled as instantaneous synapses that increase the membrane potential of the afferent neuron by 25mV, while inhibitory connections decrease it by 25mV, mirroring the excitatory synapses (Supplemental Table S3). The thalamocortical part of the model is adapted from [23] and is defined below.

### 2.5 Conductance-based Model for cortical neurons

Excitatory and inhibitory (E and I) cortical neurons were modeled as conductance-based neurons, adapted from [23]. The membrane potential 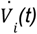 of the neuron *i* at time *t* is described by the following equation:

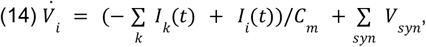

where *C*_*m*_ = 1µ*F*/*cm* denotes the membrane capacitance, and 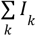 is the sum of ionic currents that pass through ion channel *k*. Unlike the receptors, TG and BS neurons, for the thalamic and cortical neurons, the full action potential is modeled. Synapses are modeled as instantaneous synapses: when a presynaptic neuron spikes, the postsynaptic membrane potential is increased by an amount *V*_*syn*_, whose value is described in Supplemental Table S3. *I*_*i*_ (*t*) corresponds to an intrinsic noise current of neuron *i* behaving as

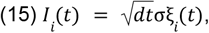

where the timestep of the simulation for cortical neurons is dt = 0.01ms, ξ_*i*_ is a random value drawn at every step for each neuron independently from a normal distribution centered around *μ* = 0 and with standard deviation 1, which is then multiplied by the amplitude 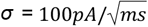.

The ionic currents *I*_*k*_ used in equation (14) for excitatory (E) and inhibitory (I) neurons are adapted from the work by Pospischil and colleagues from 2008 [24]:

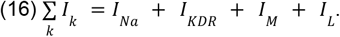

The membrane potential is depolarized by a sodium current (Na) and repolarized by a delayed rectifier potassium current (KDR). Hyperpolarisation is achieved through a slow non-inactivating potassium current (M) and the leak current (L) maintains the neuron’s membrane potential at its baseline in the absence of input.

The sodium current is described by

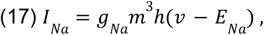

Where *g*_*Na*_is the maximum channel conductance, *E*_*Na*_ is the reversal potential of the sodium channel (Supplemental Table S2), *m* and *n* are, respectively, the activation and inactivation variables of the channel such that:

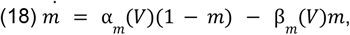

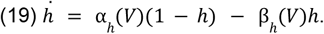

Here α is the transition rate from open to closed, and β is the transition from closed to open. Both α and β depend on a fixed threshold *V* (Supplemental Table S2) as described below:

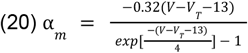

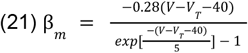

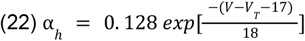

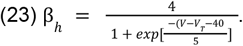

The delayed-rectifier potassium channel’s behavior is described as

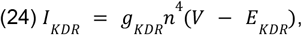

Where *g*_*KDR*_ is the maximum channel conductance, *E*_*KDR*_ is the reversal potential (Supplemental Table S2), and *n* is the activation is the channel described by

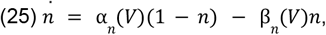

with the transition rates α andβ given by:

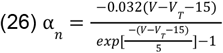

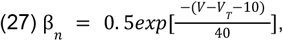

The slow non-inactivating potassium current *I*_*M*_ is described by

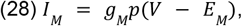

Where *g*_*M*_ is the maximum channel conductance, *E*_*M*_ is the reversal potential (Supplemental Table S2), and *p* is the activation variable of the channel described by

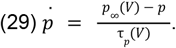

Here *p*_∞_ is the steady-state activation function defined as:

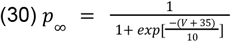

and *τ*_*p*_ is the channel time constant depending on *τ*_*max*_ (Supplemental Table S2) as:

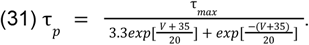

Finally, the leak channel current is described by

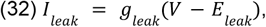

With *g*_*leak*_the maximum channel conductance and *E*_*leak*_ the reversal potential of the channel (Supplemental Table S2).

### 2.6 Parameter Fitting for Cortical Neurons

The distinction between E and I neurons in this model was made through channel-specific parameters to mimic known differences between their input-output characteristics [24]. E neurons typically emit action potentials at a lower frequency than I neurons in the barrel field of the primary somatosensory cortex when stimulated with injected current with a similar amplitude [25]. The *g*_*Na*_, *g*_*M*_, *g*_*leak*_, *V*_*T*_ and *τ*_*max*_ from the neuron model described above were fitted to reproduce the current-frequency (I-f) curves from Zeldenrust et al. (2024) [25].

### 2.7 Model of Thalamic Neurons

Thalamocortical relay (TCR) neurons are also modeled as single-compartment models and are based on a model from Ching et al. (2010) [26]. For these neurons, the membrane potential *V* of the neuron *i* over time *t* is described as follows:

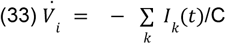

Following the description of the cortical neuron model 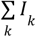 is the sum of ionic currents passing through the voltage-gated channels *k*. For every spike originating from presynaptic neurons in the brainstem (BS), the postsynaptic TCR cell’s membrane potential was increased by 10 mV as to spike when receiving a spike. In this model, following the TCR model from Ching et al. (2010) [26], the ionic currents can be expressed as:

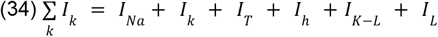

The sodium channel (Na) dynamics are the same as for the cortical neurons, described in equations (17) to (23), with *g*_*Na*_ = 90*mS*, and *E*_*Na*_ = 50*mV* (Supplemental Table S2).

The potassium (K) current is described by

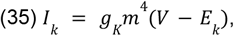

with the maximum conductance *g*_*K*_ = 10*mS*, and the reversal potential *E*_*K*_ = −100*mV* and the channel activation variable *m* given by

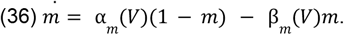

Here, transition rates α and β are given by

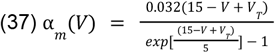

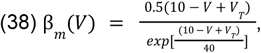

with threshold *V*_*T*_ = 25*mV*.

In addition to the sodium and potassium currents, TCR neurons exhibit a T-type calcium current

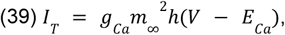

where *g*_*Ca*_ = 2*mS* is the maximum channel conductance, *E*_*Ca*_ = 120*mV* is the reversal potential, and *m* represents the channel activation based on the steady state given by

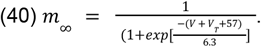

Similarly, *h* represents the inactivation based on its steady state *h*_∞_

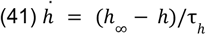

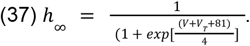

The time constant *τ* is described as:

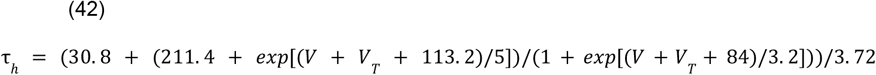

Both *m* and *h* depend on a fixed threshold *V*_*T*_ = 2*mV*.

The TCR neurons also have an h-current, of which the dynamics are described by

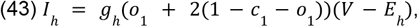

where *g*_*h*_ = 0. 01 *mS*/*cm*^2^ is the maximum conductance of the channel, *E*_*h*_ =−40*mV* the reversal potential and *o*_1_ and *c*_1_, respectively, the channel activation and inactivation variables described by:

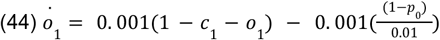

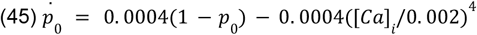

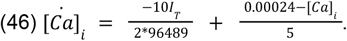

Here, *p*_0_ modulates the h-current according to equation (45), based on the calcium concentration in the cell (equation 46).

The inactivation *c*_1_ is given by

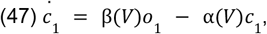

with transition rates α and β given by

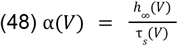

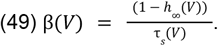

Here *h*_∞_ is the steady state value and *τ*_*s*_ the time constant such that:

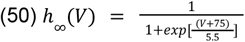

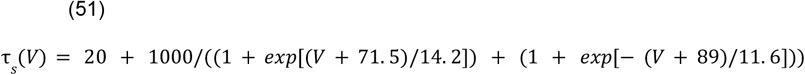

TCR neurons contain two leak channels (potassium: *I*_*K*−*L*_, and general *I*_*L*_) described by

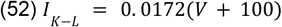

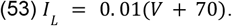

### 2.8 Parameter fitting for Thalamic Neurons

The initial TCR neurons model from Ching et al. (2010) [26] modeled the thalamic neurons during anesthesia. This state makes the TCR neurons burst upon stimulation. To reproduce thalamic activity in an awake state, we depolarized the membrane potentials of the TCR neurons by increasing the maximal conductance of the hyperpolarization-activated h-current *g*_*h*_ (Supplemental Table S3).

### 2.9 Network Architecture

The network was composed of 12 populations, among which, three input populations:

- The SA1 population was composed of 24 neurons
- The SA2 population was composed of 10 neurons
- The RA population was composed of 36 neurons
- TG, BS, and VPM populations were composed of 70 neurons
- The L4E population was composed of 188 neurons
- The L4I population was composed of 33 neurons
- The L3E population was composed of 158 neurons
- The L3I population was composed of 28 neurons
- The L2E population was composed of 86 neurons
- The L2I population was composed of 15 neurons

The connectivity between the neuron populations is described in Supplemental Table S3. Simply, all populations up to BS (excluded) are connected with a one-to-one connectivity pattern following the scheme in Figure 1d. Once within BS, the connections become probabilistic. For instance, a BS neuron has a 1% probability of sending a lateral inhibitory connection to another BS neuron, which means that amongst (70^2^ − 70) all possible connections among all BS neurons, 1% of them will exist. BS neurons are both excitatory (feedforward connections to VPM) and inhibitory (lateral connections to BS) [27]. VPM contains only excitatory neurons. Starting from cortical L4, each structure contains excitatory and inhibitory populations, which have their own connectivity patterns as described in Supplemental Table S4.

### 2.10 Information Quantification

We quantified the information transferred across layers within the modelled pathway by computing the mutual information between the stimulus and the firing rate of a whole population or of a single neuron for each stimulus. This MI was computed as follows:

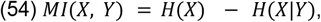

with *H* being the Shannon entropy, X the distribution of spikes for a given population or a given neuron over trials, and Y the distribution of the stimulus conditions over trials. No bias correction was applied to the mutual information (MI) calculation. Although this might seem unexpected for biological data, our analysis concerns a deterministic model: for a given stimulus, the neural response is always identical. Consequently, subsampling bias cannot occur. The absence of noise ensures a single possible output for each stimulus, though distinct stimuli can converge onto the same neural representation. In such cases, the conditional input entropy given the output, H(X∣Y), may be smaller than the input entropy H(X), since two stimuli x and x′ can produce the same output y. Nevertheless, the mapping remains deterministic: whenever x or x′ is presented, the same response y will occur. For these reasons, bias correction was not required in our MI estimates.

The resulting MI was plotted against the entropy of the stimulus computed as:

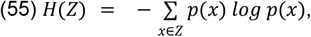

where Z is the distribution over trials of either the stimulus values, the firing rate of a single neuron, or the firing rate of a whole population.

Based on the MI values computed for each neuron, we identified the most informative neuron for each population for each condition (Figure 2) and used its firing rate as input for decoders (see Decoders)

### 2.11 Quantification of neural correlations

To determine the redundancy within each structure, we computed the Pearson correlation coefficient within each population across pairs of neurons’ spike counts. It is given by

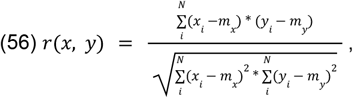

with *m*_*x*_ the mean of vector *x*_*i*_ and *m*_*y*_ the mean of vector *y*_*i*_, with *x*_*i*_, the firing rate of neuron x in response to stimulus *i*, and *y*_*i*_, the firing rate of neuron y in response to stimulus *i*. N is the number of stimuli presented to the network for a given parameter, modified from the baseline.

### 2.12 Decoders

For each input class (whisker or finger), we picked three decoders and trained them to retrieve the stimulus parameters from the neural activity, from a single population or single neuron. For instance, if the changed parameter was the amplitude of deflection (in the whisker case), the decoder had to classify the stimulus amongst the 19 possible amplitude values. For fingertip and whisker data, the decoders relied on a vector containing the number of spikes for each neuron of a population as an input for a given trial. For a population of size P, each trial had an input of size 1xP and produced a single value. In single-neuron decoding, the input was a scalar representing the number of spikes produced in a given trial. From scikit-learn [28], we used 3 types of classifiers: RidgeClassifier, NaiveBayes, and MLPClassifier. We used the accuracy of the classification as a metric to measure the performance of the decoder, described as follows:

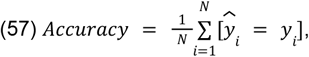

with *N* the number of trials, 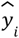 the predicted label of the trial *i* and *y* the true label, and *Accuracy* ϵ [0; 1], where 1 represents perfect prediction.

For the fingertip data, we tasked the decoders to perform a regression on the 160 different amplitudes applied during the psychophysiological task. We relied on the Ridge, BayesianRidge, and MLPRegressor functions from the scikit-learn library [28]. The decoders’ performance was assessed through *R*^2^ as follows:

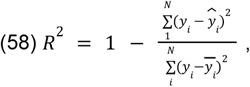

where *y* and 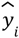 denote the true and predicted values, respectively, 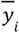 is the mean of the observed values. A value *R*^2^ = 1 indicates perfect prediction, *R*^2^ = 0 indicates performance equivalent to predicting the mean of the data, and negative values indicate worse-than-mean prediction.

## Results

To examine stimulus representation and information flow in the tactile system, we created a model of the ascending somatosensory pathway. We extracted spiking activity generated in each layer after stimulation of the input layer by output from either a virtual whisker ([16], see section 2.1.1) or from recorded skin displacement at the fingertip in a psychophysics experiment (see section 2.1.2).

We first verified that activity from the mechanoreceptors propagated through the entire model up to the last layer of the cortex. We then quantified the transferred information between the stimulus and the resulting single-neuron or population response at each stage of the pathway. We further analyzed the coding structure through pairwise correlation among neurons’ activity within each population. Finally, we assessed the quality of stimulus representation through the ability of decoders to recognize tactile stimulus parameters from spiking activity (single-neuron and population).

### 3.1 Model Description

We fitted the parameters of an AdeX neuron [21] to the instantaneous firing rate recorded by Sonekatsu and Gu in 2019 [19] (see Methods and Supplemental Table S1). Through this procedure, we obtained three input populations that responded like the recorded mechanoreceptors in the whisker follicle to ramp-and-hold stimuli. The TG and BS neurons were tuned to spike whenever they received synaptic input. The cortical excitatory neurons were tuned to have regular spiking patterns, while the inhibitory neurons were tuned to have fast spiking characteristics [23].

The result is a neural network model that describes the propagation of neural activity from the touch cells up to the cortex upon whisker stimulation (Figure 1). The activity profile remains the same from TG to the layer 2 of the somatosensory cortex: It shows a sharp surge of activity at the onset of a stimulus, a decrease toward a plateau phase during the hold part of a ramp-and-hold, and another increase of activity when the force is released (Figure 1). A clear sparsification of activity is noticeable in the layers 2 and 3. Upon the application of a force on the 3D virtual whisker, activity appears in the receptor population and propagates across each structure up to layer 2 of the somatosensory cortex (Figure 3). A shift in the stimulus, either in amplitude of deflection, direction, frequency, speed, or the pole position along the whisker results in a shift in the neural response for every population, easily noticeable in the periphery (Figure 3 and supplemental Figures 1 to 4) except for SA1, which appears unaffected by speed or frequency of the stimulus or SA2, which was not triggered during the frequency paradigm as the force applied to the whisker was not sufficient to trigger activity in these receptors (supplemental Figure 3). Consistent with whisker-related input, when presented with skin displacement recordings, the activity propagated in the entire network, and different amplitudes of displacement resulted in different neural activity (supplemental Figure S17).

**Figure 3.**
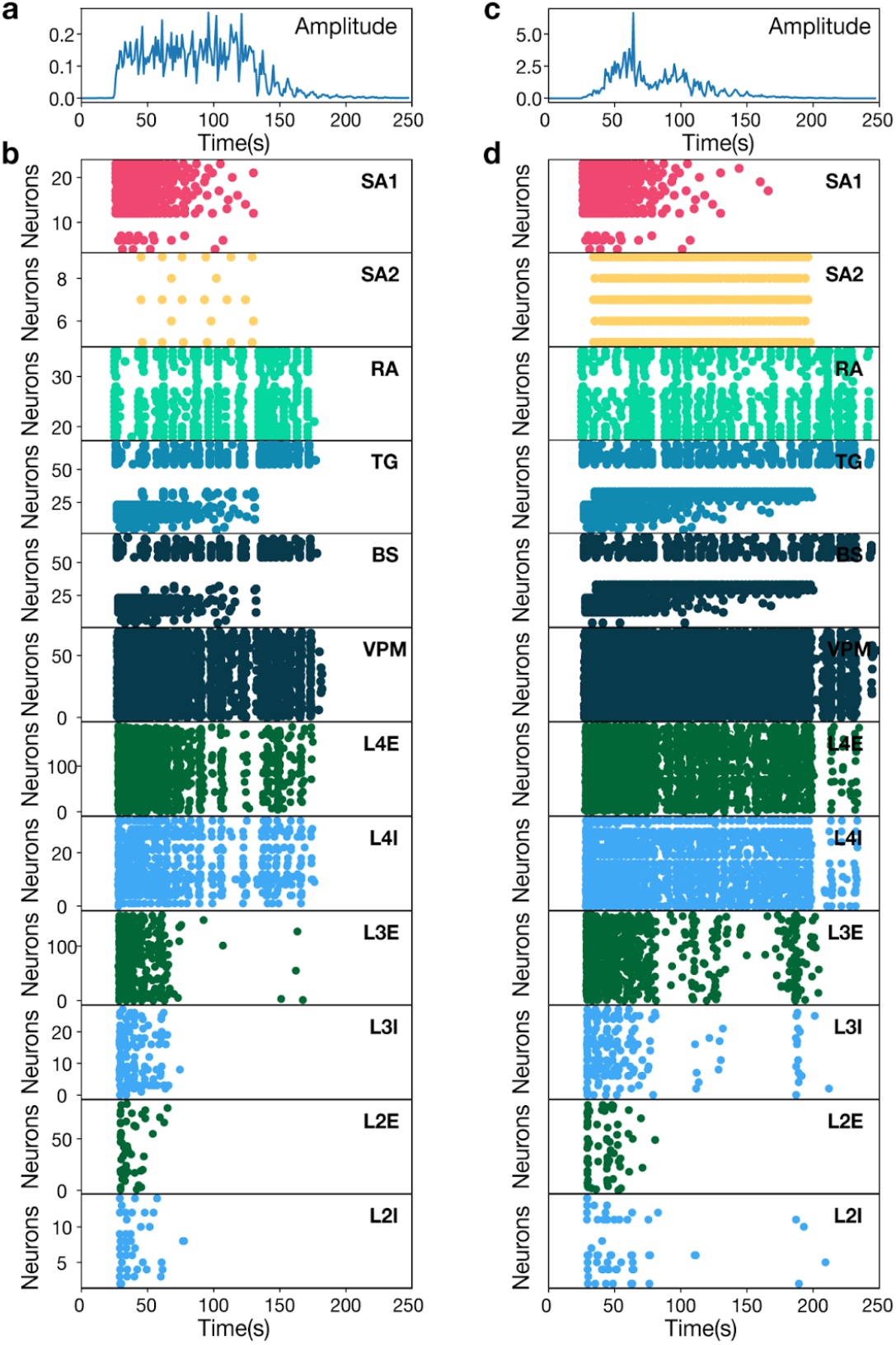
Example model responses to a ramp-and-hold whisker stimulus to two different amplitudes (Figure 2b). **(a)** Amplitude of the whisker displacement (vertical axis) in time (horizontal axis). **(b)** Single neuron responses (dots across the vertical axis) in time (horizontal axis) triggered by the stimulus from a, color-coded based on the population identity. From top to bottom, populations are: SA1, red; SA2, yellow; RA, light green; VPM, pastel blue; BS, dark blue; VPM, dark blue; L4E, dark green; L4I, light blue; L3E, dark green; L3I, light blue; L2E, dark green; L2I, light blue. **(c)** Same as in a, but a different pattern of the whisker displacement amplitude. **(d)** Same as in b, but responses are triggered by amplitude in c.

Overall, our network exhibits responses mimicking biological responses to a ramp-and-hold stimulus and responds to deflections of the virtual whisker from WhiskIt [16], as well as to recorded skin displacements.

### 3.2 Information transfer is maximized in the thalamus

We quantified stimulus representation at the single neuron and population level by computing the mutual information between the resulting spike counts and the range of the different stimulus parameters (for instance direction or speed).

For the whisker input, considering the periphery, we can see a distinct tuning profile (Figure 4). The activity of the SA1 population is very informative about the stimulus direction, but not so much about the other stimulus parameters; the SA2 activity doesn’t reflect differences in frequency, but is informative about the other stimulus parameters. Information transferred by the RA population is constant across tactile features. These observations are true for single neuron and population activity. We also notice the widespread MI values among single neurons. Upon entering the central nervous system, this specialization disappears, as all populations bear information on each feature and, the distribution of MI values within single neurons become notably narrower in the thalamus and the successive layers. Interestingly, information within single neurons consistently drops in L3 and L2 when compared to L4. Similarly, we notice a slight decrease in information at the population level starting in layer 3, but in particular in the L2E population, compared to L4.

**Figure 4.**
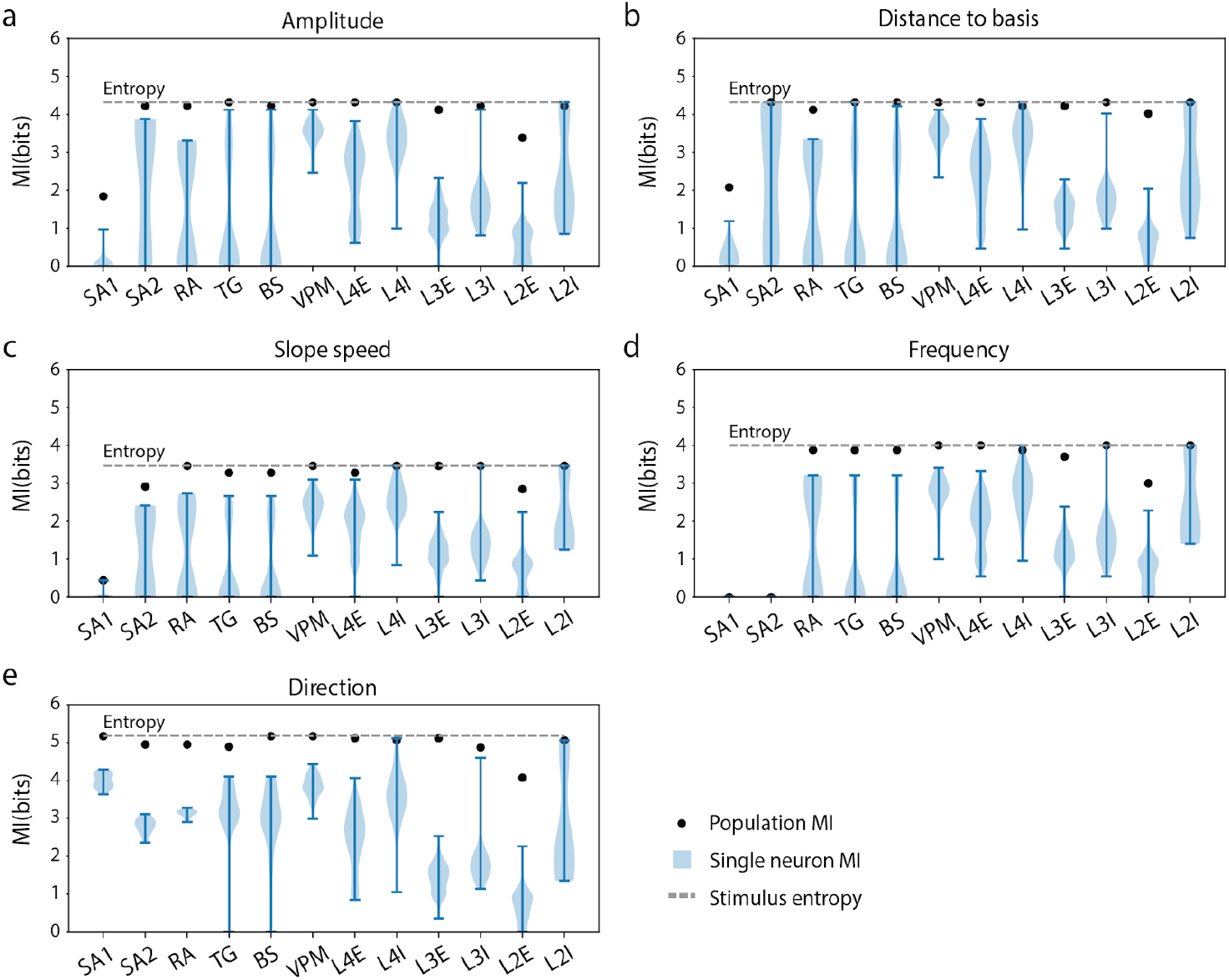
Mutual information (MI) distribution over the populations from simulated whisker displacements. **(a)** The mutual information (MI) calculation between the firing rate of each neuron within a population and the stimulus amplitude, in an experiment where the amplitude was varied (see Figure 2.b) (violin plots; legend label: Single neuron MI), population information rate (dot; legend label: Population MI), and the entropy of the stimulus (dashed line; legend label: Stimulus entropy) is shown for each population (horizontal axis) **(b)** Same as in a, but for stimulus distance variation, see Figure 2.c). **(c)** Same as in a, but for stimulus speed variation, see Figure 2.d). **(d)** Same as in a, but for stimulus frequency variation, see Figure 2.e). **(e)** Same as in a, but for stimulus direction variation, see Figure 2.f).

For the skin displacement recording input, single neuron activity conveyed substantially less information than population activity about the stimulus amplitude (supplemental Figure S14a). However, the observation made on the MI using the whisker input holds when fingertip data is used. Single-neuron’s MI distribution is broad up to BS and narrows down starting from the thalamus. MI at the population level is maximised in VPM and the L4 and decreases drastically in the L2 excitatory population, in the successive layers. In contrast to whisker input, RA’s single-neuron MI distribution is narrow.

These results demonstrate the capacity of the model to propagate tactile information from all mechanical features of stimuli from different systems, while revealing convergence and compression of coding as signals ascend to the cortex.

### 3.3 Neural activity is decorrelated in the central nervous system compared to the periphery

We assessed the similarity of the coding structure across neurons by computing the Pearson correlation coefficient between spike counts of individual neurons within each population.

As shown in Figure 5, neuronal activity in the periphery was highly correlated between neurons across all tactile features. The exception was the frequency paradigm: there, SA1 and SA2 neurons exhibited no spikes, resulting in an artificially null correlation. Amongst the mechanosensitive cells, both SAs populations displayed the highest within-population correlations. Upon entry into the brainstem, correlations between neurons of the same population decreased and remained stable in the successive layers.

**Figure 5.**
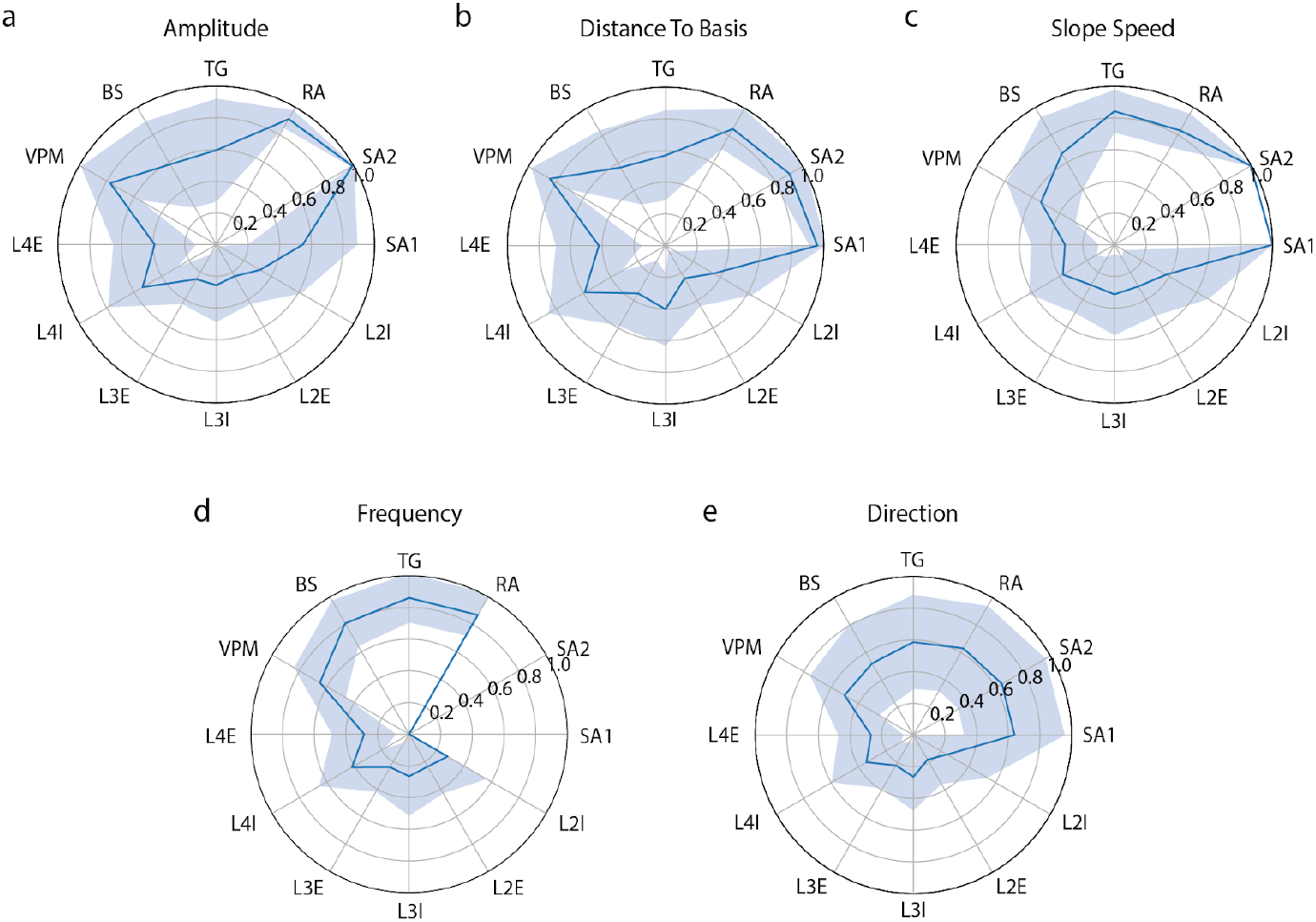
Correlations between single neuron activities in different populations. **(a)** Pairwise Pearson correlation between the firing rates of each neuron pairs within a population in response to a shift in deflection amplitude (fig. 2.b). The solid line denotes the mean across all pairwise correlations, and the shaded region the mean ± the standard deviation. **(b)** Same as in a, but for a shift in distance (fig 2.c) **(c)** (Same as in a, but for a shift in speed (fig 2.d) **(d)** Same as in a, but for a shift in frequency (fig 2.e) **(e)** Same as in a, but for a shift in direction (fig 2.f)

In contrast, individual neuron activity resulting from skin displacement at the fingertip, while keeping a high correlation in the periphery, showed an increase in correlation in the thalamus (supplemental Figure S14.b). Notably, the correlation still decreased in the previous layer (BS). Nonetheless, the lowest correlation was achieved in layer 4 of the cortex and remained stable in the downstream layers.

These results demonstrate a reduction in redundancy among neurons as tactile information ascends to the cortex, reflecting a diversification of coding strategy in the central nervous system.

### 3.4 Stimulus decoding

We evaluated the model’s ability to represent tactile stimulus features by training decoders on spike counts at the population and single-neuron levels. For the whisker input, we performed classification, and for the fingertip skin-displacement input, we performed regression (see Method 2.12). In both cases, decoders were trained first on a vector of spike counts per neuron within a population and then on a scalar containing the spike count of the most informative neuron within that population.

In accordance with the MI results (section 3.2), the decoders were capable of retrieving the parameters of the applied stimuli from the spike counts per neuron for each population and stimulus condition (Figure 6). With the whisker input, the exception is SA1, which provides a reliable readout only for direction decoding. Notably, MLP was outperformed by linear and naive Bayesian decoders when trained on the peripheral populations (TG, SA1, SA2, and RA). In contrast, fingertip stimuli appeared rather difficult to distinguish based on neural activity. Keeping in consideration that the task was a regression, we see that MLP outperformed the other decoders, no matter which population’s readout was used, and was the only algorithm capable of decoding the stimulus feature accurately from VPM and the inhibitory populations within the cortex. Bayesian and linear regressors achieved a similar performance, but lacked accuracy when relying on inhibitory populations. For each decoder, decoders trained on RA and SA1 activity couldn’t provide a reliable readout.

**Figure 6.**
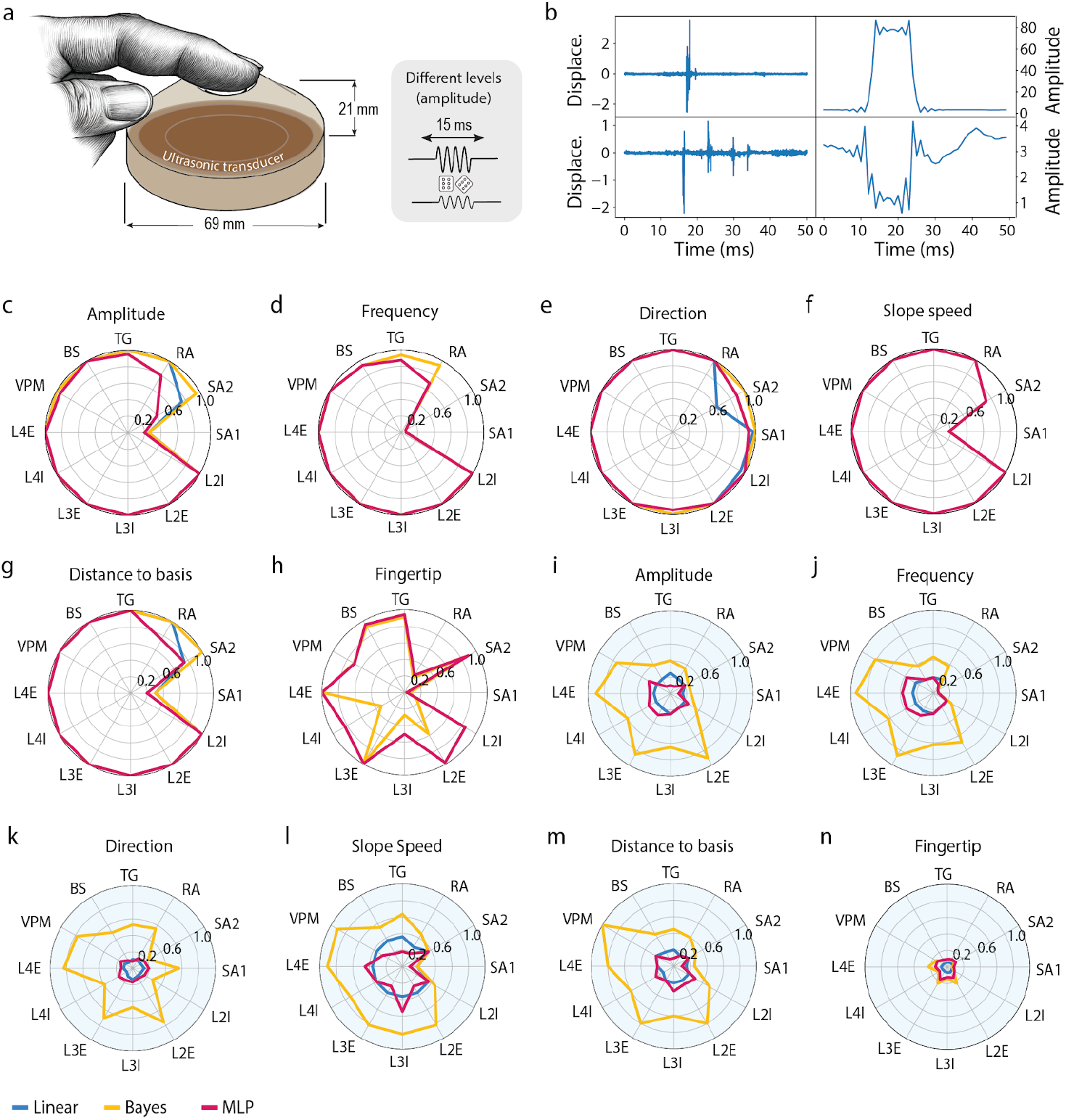
Decoding accuracy of the different populations. **(a)** Experimental setup of the sonication experiment. An ultrasonic transducer sends waves to the fingertip. **(b)** Two different amplitudes of skin displacement. Left: Skin displacement (Displace.) recording. Right: Pre-processed input from left. **(c)** Polar plots representing the accuracy ϵ [0;1] of decoders (legend below the panel k: blue, linear; yellow, Bayesian; red, MLP) trained to retrieve the amplitude of the tactile input. The decoding was done based on the spike count per neuron for an entire population. **(d)** Same as in c, but for decoding frequency. **(e)** Same as in c, but for decoding direction. **(f)** Same as in c, but for decoding speed. **(g)** Same as in c, but for decoding distance. **(h)** Same as in c, but for decoding skin displacement on a fingertip. Fingertip values are based on *R*^2^ ϵ [−∞; 1]. **(i)** The polar plot represents the accuracy ϵ [0;1] of decoders (legend below the panel k: blue, linear; yellow, Bayesian; red, MLP) trained to retrieve the amplitude of the tactile input. The decoding was done based on the spike count on the single most informative neuron per population. **(j)** Same as in i, but for decoding frequency. **(k)** Same as in i, but for decoding direction. **(l)** Same as in i, but for decoding speed. **(m)** Same as in i, but for decoding distance. **(n)** Same as in i, but for decoding skin displacement on a fingertip. Fingertip values are based on *R*^2^ ϵ [−∞; 1].

When tasked to decode whisker stimulus features from the single most informative neuron within each population, only the naive Bayesian classifier could learn, achieving near-perfect accuracy relying on VPM, L4E, and L3E read-outs (Figure 6). Overall, this decoder exceeded 0.5 accuracy with activity from all structures downstream from BS (excluded). In the fingertip scenario, the read-out from a single neuron was not sufficient to perform a regression between the amplitude of skin displacement and the spike count.

These results demonstrate a robust representation of tactile features within the different layers of our model. They show that single-neuron activity, starting from VPM, is sufficient to perform a classification task, but falls short of results when asked to perform a regression.

## Discussion

We introduce a spiking neural network model that reproduces the ascending somatosensory pathway from mechanoreceptors to cortex and demonstrate its ability to integrate tactile information spanning all mechanical features of tactile stimuli. By applying inputs derived from both virtual whisker deflections and recorded fingertip skin displacements, we show that the model can respond to a broad range of stimuli. We confirm that population activity at each stage of the pathway retains sufficient information to identify a stimulus through information-theoretic analyses and decoding performance. We furthermore demonstrate that in the case of whisker input, single-neuron activity in the thalamic and cortical layers is reliable enough to train the decoders, as long as the most informative neuron is used. Together, these findings indicate that hierarchical organization and convergent projections in the ascending pathway result naturally in efficient, low-redundancy, and robust population codes. This architecture follows neuromorphic designs principles (sparse spiking, layered processing) and can be adapted for deployment on neuromorphic hardware.

### 4.1 Stimulus propagation across the network

Our results demonstrate the specialization of the periphery through the various responses of the mechanoreceptors. Indeed, when we take a look at the diverse dimensions of tactile stimuli we explored, we notice a discrepancy between the information transferred across the mechanosensitive cells. For instance, while every receptor is sensitive to direction, only the RA population was found to be informative about frequency. Regarding the other dimensions, the SA2 and RA populations were both highly informative, whereas the stimulus information in the SA1 population only peaked for direction (Figure 4). This diversity in stimulus representation within the peripheral system allows our model to transmit information spanning the entire range of tactile features. Upon reaching TG, this specialization disappears. It is known that peripheral sensory cells convey partial information on tactile stimuli [2, 6]. The convergence of these multiple cells’ outputs within a single structure explains the collapse of this specialization within TG. Similar to our model, in biology, while SAs and RAs cells remain distinguishable in BS [29], it appears that the converging SA1 and SA2 connections into a single post-synaptic neuron would result in a similar phenomenon. Furthermore, the activity patterns demonstrated by our model follow the stereotypical responses seen in biology, composed of a highly synchronized population response at the onset and offset of the stimulus and a lower plateau phase during the stimulus presentation (Figures 1 and 2). This pattern is maintained throughout the entire ascending somatosensory pathway, regardless of whether virtual whisker input or measured skin displacement at the fingertip was used as input. The conservation of this pattern in the cortical area is especially important, as the high synchronicity of neural responses at the onset of the stimulus is believed to be critical to trigger the cortical damping effect [30]. This allows a limitation of excitatory neurons’ activity by their inhibitory counterpart, thus allowing cortical neurons to be more sensitive to stimulus timing rather than the amplitude of thalamic neurons’ response. In the context of the whisker system, it is especially important to distinguish between the stimulation of different whiskers. This phenomenon is reproduced in our model.

### 4.2 Decoding performances on population activity

The propagation of a stimulus representation across the entire ascending somatosensory pathway, across all the tactile features that were presented, allows us to train decoders on the resulting activity of the different layers. When relying on the entire population activity, that is, the total number of spikes produced within a population for a given trial, even the simplest decoder (RidgeClassifier) was able to retrieve the stimulus identity from the activity of most populations. The exceptions belong to the periphery and remain as such across all the different decoders that were used. Furthermore, the lack of accuracy from the classifiers is aligned with the MI calculation from these populations. These results hint toward an effect of the receptors rather than a limitation of the decoding algorithms, with SA1 being informative only about the direction, and both SAs populations being non-informative about frequency. This demonstrates that even at the population level, the representation of the entire span of a tactile stimulus is only possible when relying on a diverse set of receptors [2]. The downstream neurons receive direct or indirect input from multiple classes of mechanoreceptors, allowing them to represent the entire range of tactile dimensions.

### 4.3 Information flow in the ascending somatosensory pathway

Consistently across all stimuli, we notice a pattern regarding the information flow. The periphery shows a wide distribution of MI (between single neurons’ spiking activity and the stimulus) values across neurons, while it has a narrow distribution as soon as it reaches the thalamus. When looking at the Pearson correlation across the firing rate of single neurons within a population, we see that the narrowing of MI distribution happens together with decorrelation, starting in BS. This phenomenon results in the ability of the classifiers to decode information from a single neuron within the thalamus and layers 4 and 3 of the somatosensory cortex model. Interestingly, the maximum MI values and a low decorrelation happens within the bursting thalamic population (VPM), confirming the results from Guo et al. (2021) [13], that demonstrated that bust coding reduced redundancy while preserving discriminability. Single neuron activity was provided less information when the model was presented with a more noisy stimulus (recorded skin displacement at the fingertip), thus demonstrating the robustness of population coding as opposed to single neuron coding. With this in mind, the central nervous system appears to decrease redundancy across neurons while preserving a robust representation, thus maximizing efficiency.

### 4.4 An adaptable tool

A key outcome of this work is the model itself, which we provide as an adaptable open-source tool. Built in the Brian2 simulator with a modular structure, users can easily modify neuron equations, synaptic dynamics, or connectivity via a configuration. For instance, one could replace the AdEx mechanoreceptor models with a more complex model of mechanotransduction, or swap out the cortical neuron model for a different one. The provided analysis functions (for computing information, correlations, decoding, etc.) are general and can work with any spiking data, making it straightforward to plug in alternative network configurations or novel neuromorphic chip outputs for benchmarking.

The versatility of our model is demonstrated by its handling of two different sensor modalities (whisker vs. skin) with minimal changes. This suggests that the model can be potentially extended to any mechanosensory input (e.g., foot vibrations in rodents or touch in other species) by adjusting the first layer tuning. It “paves the way” for a unified understanding of tactile processing along the pathway by providing a sandbox to simulate injuries (lesion a layer and see effect), neurological disorders (e.g. neuropathy could be simulated by adding noise or dropping the input layer from the model), or comparing encoding strategies by modifying neuron’s dynamic (e.g. burst vs. tonic firing, as in [13].

From a neuromorphic engineering viewpoint, our model is directly relevant to designing low-power tactile sensing systems. It suggests that a multi-layer event-driven architecture, where each layer performs specific transformations (e.g., feature integration, redundancy reduction), can achieve efficient encoding. Neuromorphic chips could implement similar layered processing, e.g. initial silicon “receptors” that spike upon touch, a second layer that sparsifies, a third that does something akin to thalamic relay. Our results emphasize that maintaining timing (event-driven onset signals) and including inhibition for decorrelation are important design principles. Moreover, our model achieves good performance with relatively few spikes (sparse firing except at onset), which implies it would be very power-efficient on hardware that only computes on spikes (since most neurons are quiet much of the time). This aligns with the broader goal of neuromorphic tactile systems: to use sparse asynchronous events instead of dense sampling, thereby cutting down energy usage.

In conclusion, our end-to-end tactile pathway model brings together mechanistic understanding from neuroscience and practical considerations from engineering. It not only matches known biological phenomena (like adaptation, bursting, feed-forward inhibition) but also quantifies their contribution to encoding efficiency. By making this model available and modular, we hope it serves as a platform for future research – both to test scientific hypotheses about sensorimotor processing and to guide the development of advanced tactile processing algorithms in robotics and prosthetics.

## Conclusion

We have presented an end-to-end spiking neural network model of the ascending tactile pathway, from mechanoreceptors in the skin or whisker follicle to the thalamus and multi-layer cortex. The model is fitted to electrophysiological data at the periphery and built using the Brian2 simulator [15], allowing straightforward updates to neuron equations and synaptic dynamics as needed. Our model integrates all major anatomical structures of the somatosensory pathway and can be driven by both real and virtual tactile inputs, demonstrating cross-species applicability (we successfully simulated rodent whisker deflections and human fingertip displacements through the same network).

Critically, we showed that each processing stage (receptors, TG, brainstem, thalamus, cortical layers) transmits sufficient information to allow downstream decoders to identify tactile stimuli. Even the simplest decoders (linear or naive Bayesian) could often decode stimulus identity from the population spike activity at a given layer. In the case of low-noise, high-salience stimuli (e.g. a clear whisker deflections), the spiking activity of a single neuron in the thalamus or cortex was enough for a decoder to recover the stimulus feature with a high accuracy. However, for more challenging scenarios (like a graded noisy skin indentation), population-level readouts were required, highlighting the robustness of distributed coding. We also provide open-source analysis code to compute information-theoretic measures, correlations, and decoding performance on the network’s activity, illustrated with example results for both whisker and fingertip cases.

## Supplemental materials

**Supplemental Table 1.** Parameter values of the Adex model [17] for SA1, SA2, and RA receptors, as well as parameters for the elif neurons model for TG and BS.

| Parameters | SA1 | SA2 | RA | TG and BS |
| --- | --- | --- | --- | --- |
| $a$ | $20.24 \text{ nS}$ | $46.11 \text{ nS}$ | $59.1 \text{ nS}$ | |
| $b$ | $69.65 \text{ pA}$ | $3.21 \text{ pA}$ | $86.53 \text{ pA}$ | |
| $\tau$ | $44.09 \text{ ms}$ | $1882.05 \text{ ms}$ | $1890.10 \text{ ms}$ | |
| $E_{leak}$ | $- 50 \text{ mV}$ | | | |
| $V_{th}$ | $- 30 \text{ mV}$ | | | |
| $V_{reset}$ | $- 45 \text{ mV}$ | | | |
| $C_m$ | $- 0.1 \text{ nF}$ | | | |
| $g_{leak}$ | $10 \text{ nS}$ | | | |
| $\Delta_{th}$ | $5 \text{ mV}$ | | | |

**Supplemental Table 2.**
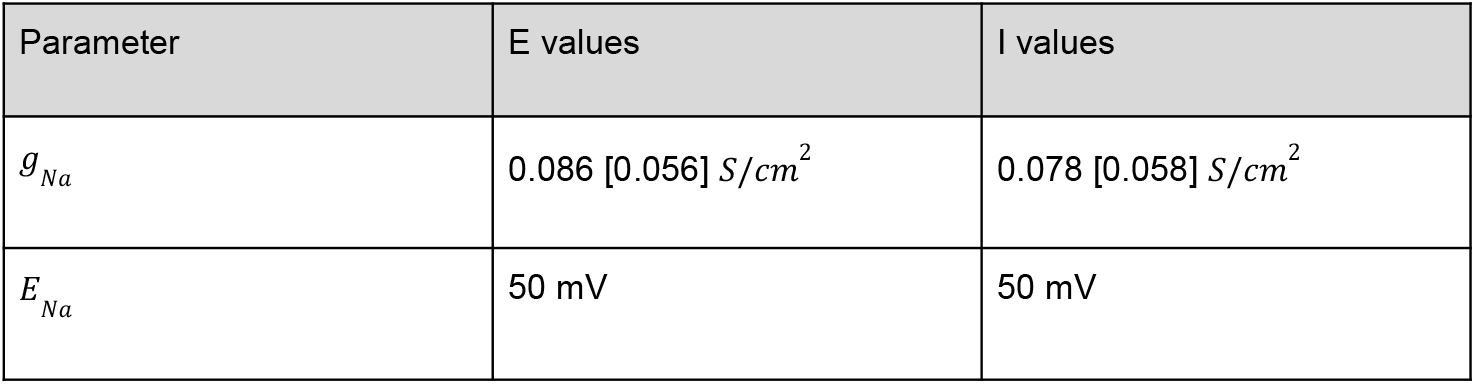
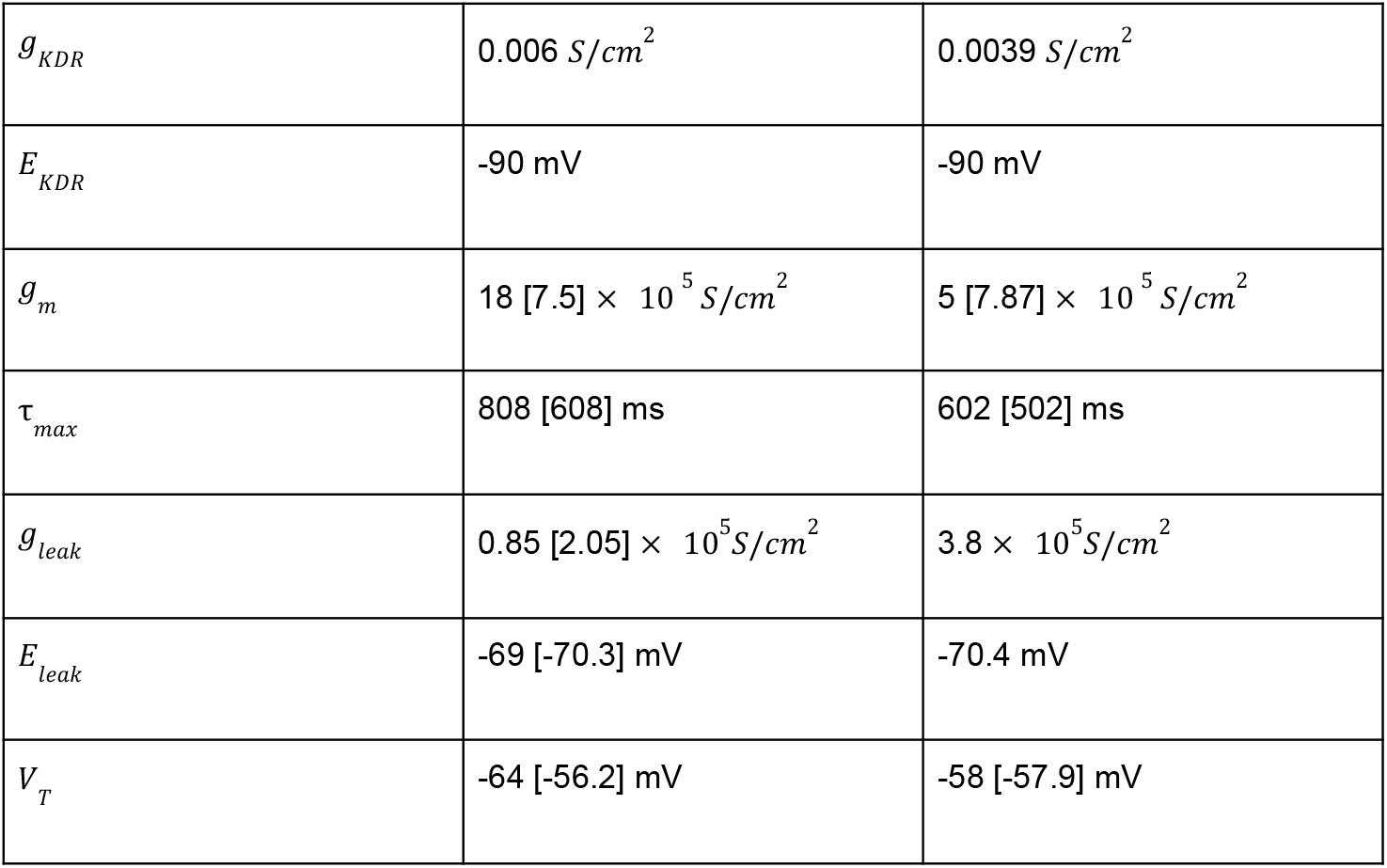
Parameter values for sodium (Na), delayed-rectifier potassium (KDR), slow non-inactivating potassium (M), and leak (L) channels used to model cortical E and I neurons. Values in brackets are default values from [19], which we changed to the outside-bracket values to adjust the neuron frequency-current behavior (method section).

| Parameter | E values | I values |
| --- | --- | --- |
| $g_{Na}$ | $0.086 [0.056] \text{ S/cm}^2$ | $0.078 [0.058] \text{ S/cm}^2$ |
| $E_{Na}$ | $50 \text{ mV}$ | $50 \text{ mV}$ |
| $g_{KDR}$ | $0.006 S/cm^2$ | $0.0039 S/cm^2$ |
| $E_{KDR}$ | -90 mV | -90 mV |
| $g_m$ | $18 [7.5] \times 10^5 S/cm^2$ | $5 [7.87] \times 10^5 S/cm^2$ |
| $\tau_{max}$ | 808 [608] ms | 602 [502] ms |
| $g_{leak}$ | $0.85 [2.05] \times 10^5 S/cm^2$ | $3.8 \times 10^5 S/cm^2$ |
| $E_{leak}$ | -69 [-70.3] mV | -70.4 mV |
| $V_T$ | -64 [-56.2] mV | -58 [-57.9] mV |

**Supplemental Table 3.** Parameter values used for simulating VPM neurons.

| Parameter | Ching et al. (2010) | Pattern-generating | Background |
| --- | --- | --- | --- |
| $g_h$ | $0.001 mS/cm^2$ | $0.01 mS/cm^2$ | $0.005 mS/cm^2$ |
| $N$ | 10 | 45 | 24 |
| $g_{In}$ | $0.05 mS/cm^2$ | $0.5 mS/cm^2$ | $0.75 mS/cm^2$ |
| $r$ | 12 Hz | 9 Hz | 5 Hz |
| $P_{In}$ | 0.5 | 0.3 | 0.1 |
| $\alpha$ | 10 | 14 | 25 |
| $\tau_r$ | 0.5 ms | 0.5 ms | 0.5 ms |
| $\tau_d$ | 2 ms | 2 ms | 2 ms |

**Supplemental Table 4.** Synapse-specific weights and connectivity patterns or connection probabilities used in the model. Note that BS neurons send lateral inhibitory connections within BS and excitatory connections to VPM; this structure doesn’t contain inhibitory neurons per se.

| Pre-post population | Pre-post neuron type | Weight (mV) | Probability to connect<br>or connectivity<br>pattern |
| --- | --- | --- | --- |
| L2-L2 | E-E | 2.8 | 5.6% |
| L3-L2 | E-I | 8.1 | 5.1% |
| L3-L2 | I-E | -17 | 5.9% |
| L3-L3 | I-I | -15 | 5.1% |
| L4-L3 | E-E | 2.4 | 5.8% |
|  | E-I | 5.7 | 0.33% |
|  | I-E | -14 | 1.7% |
| L4-L4 | E-E | 3.3 | 7.6% |
|  | E-I | 7.9 | 4.2% |
|  | I-E | -16 | 6.3% |
|  | I-I | -14 | 6.2% |
| VPM-L4 | E-E | 7 | 4.3% |
|  | E-I | 9 | 5% |
| BS-VPM |  | 10 | 30% |
| BS-BS | I-E | -50 | 1% |
| TG-BS |  | 25 | one-to-one |
| RA-TG |  | 25 | one-to-one |
| SA2-TG |  | 25 | one-to-one |
| SA1-TG |  | 25 | one-to-one |

**Supplemental Table 4.**
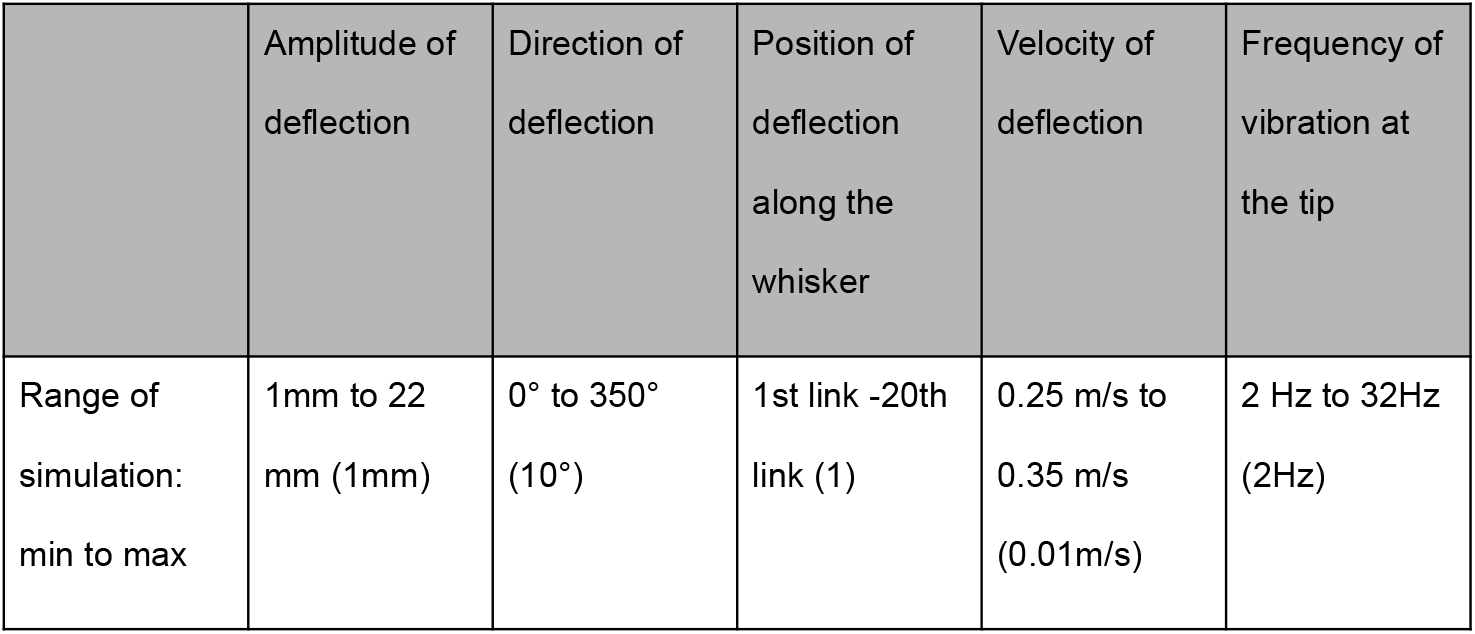
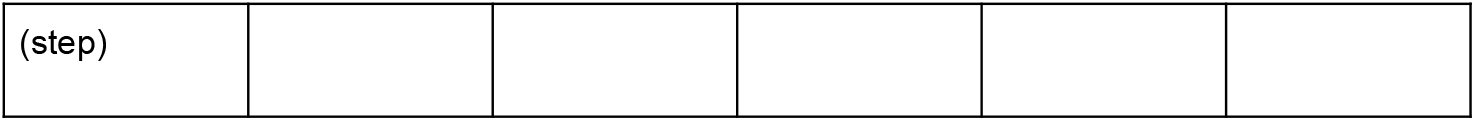
Set of parameters used for whisker deflection simulation. For each set of stimuli, the same set of initial parameters was used: The pole was positioned at 10 mm from the targeted whisker and moved at a speed of 0.3 m/s, with an angle of 0° around the x-axis (along the whisker). The amplitude of the stimulus was 1 mm, and it was held against the whisker for 100 ms at the 10th link of the whisker (middle). Then, only one of these parameters was changed, as shown in the table above.

**Supplemental Figure 1.**
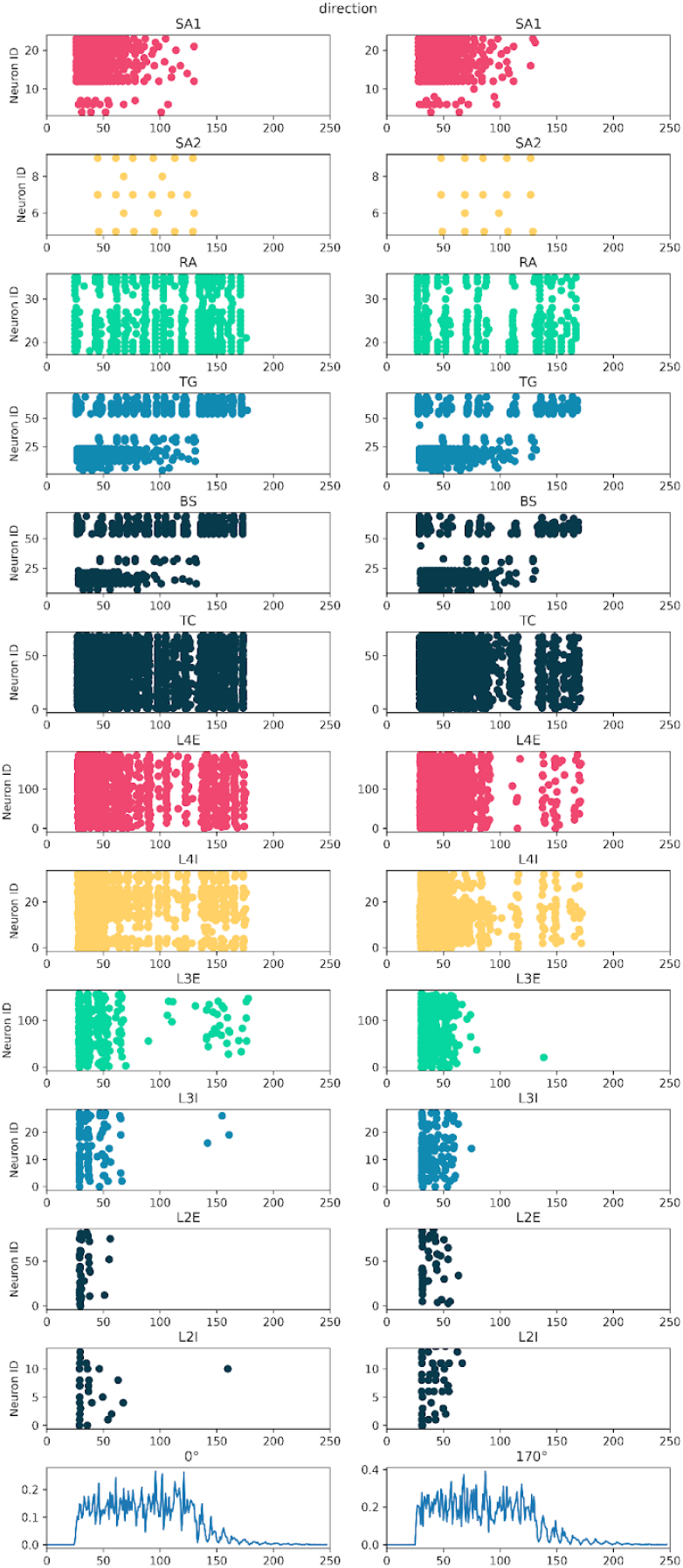
Population responses and raster plot to a direction of 0 degrees and 170 degrees

**Supplemental Figure 2.**
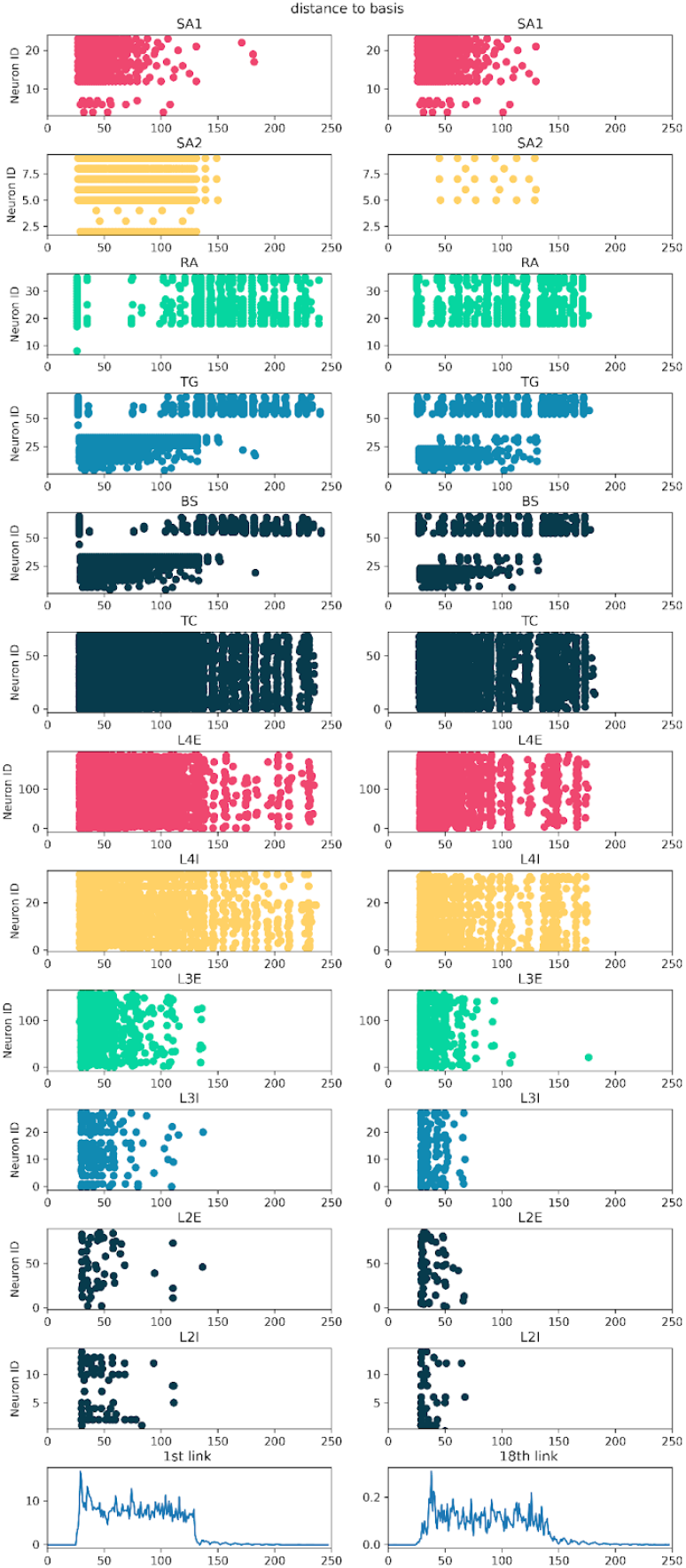
Population response and raster plot for the pole applied to the 1st link (close to the snout) of the whisker and the 18th (close to the tip) link of the whisker

**Supplemental Figure 3.**
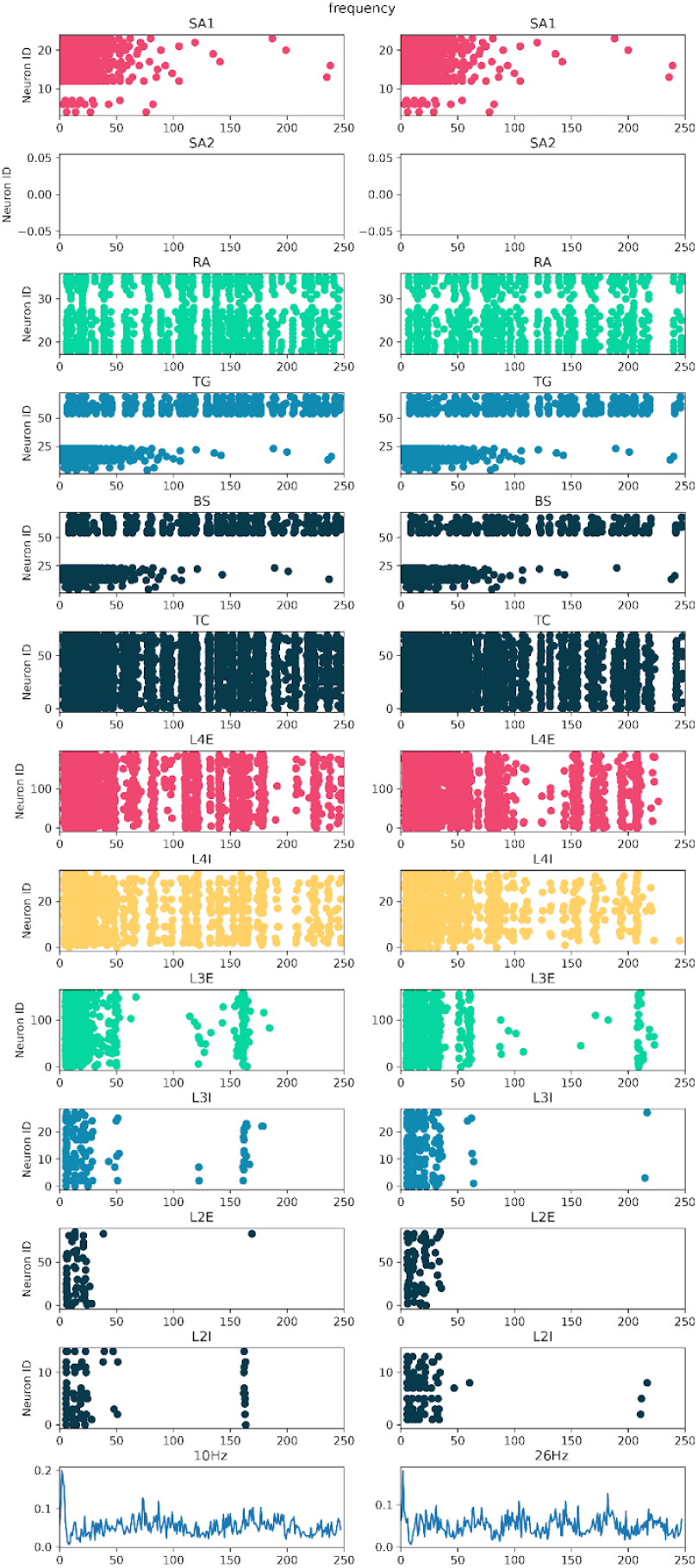
Population response and raster plot to a frequency of 10 Hz and 26 Hz

**Supplemental Figure 4.**
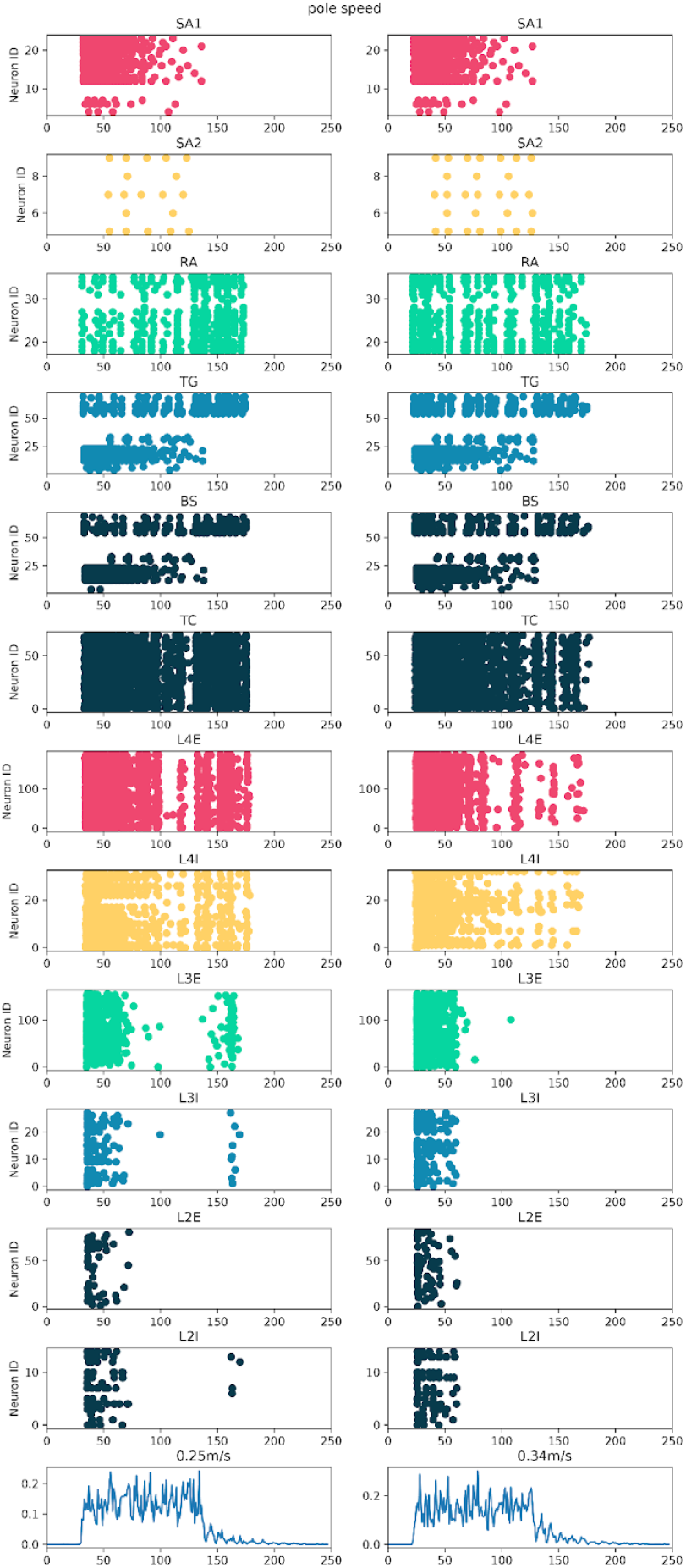
Population response and raster plot to a pole movement against the whisker of 0.25m/s and 0.34 m/s.

**Supplemental Figure 5.**
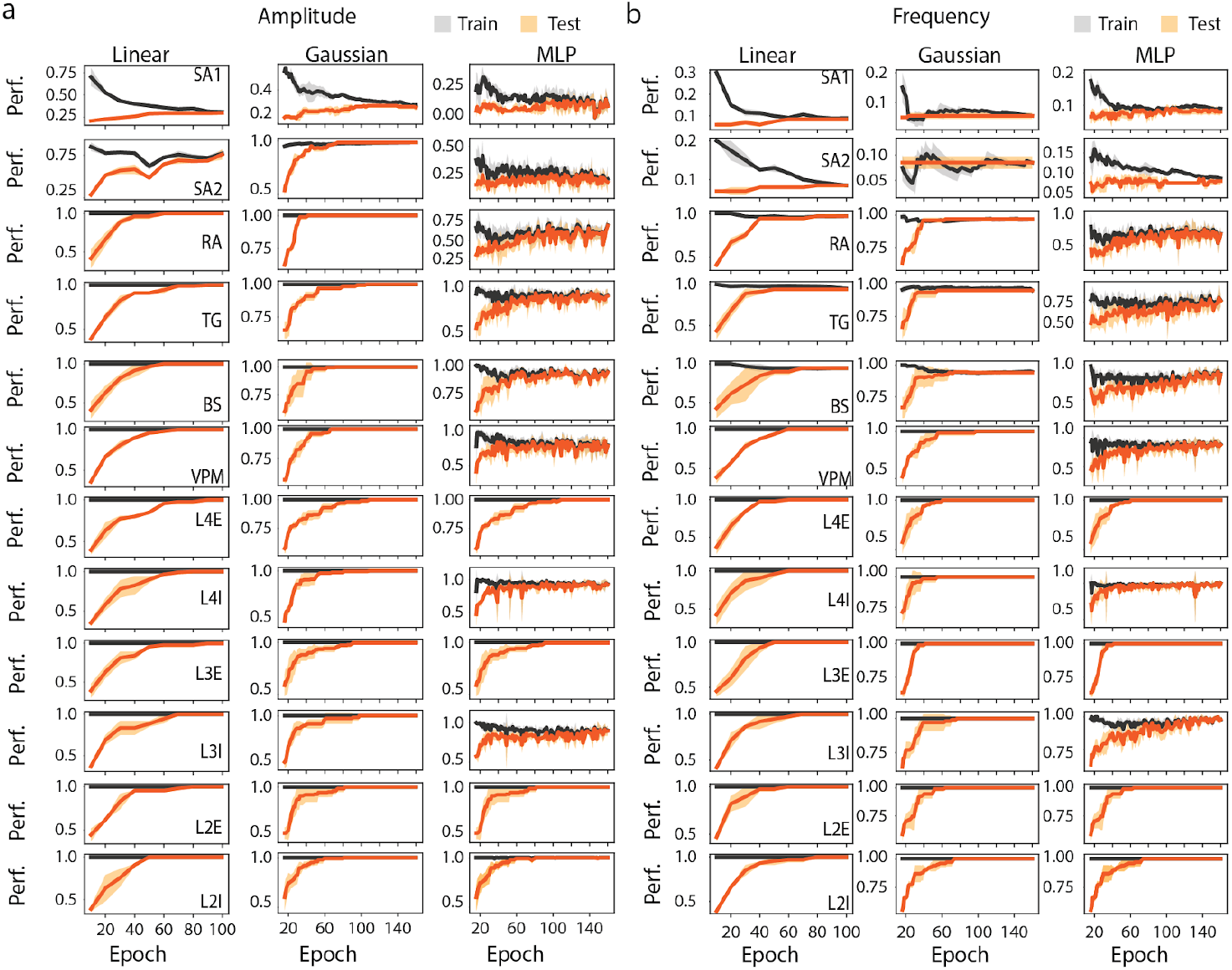
Learning curve for Linear decoder, MLP, and Bayesian when trained to retrieve stimulus **a)** amplitude and **b)** frequency, on whole population activity.

**Supplemental Figure 6.**
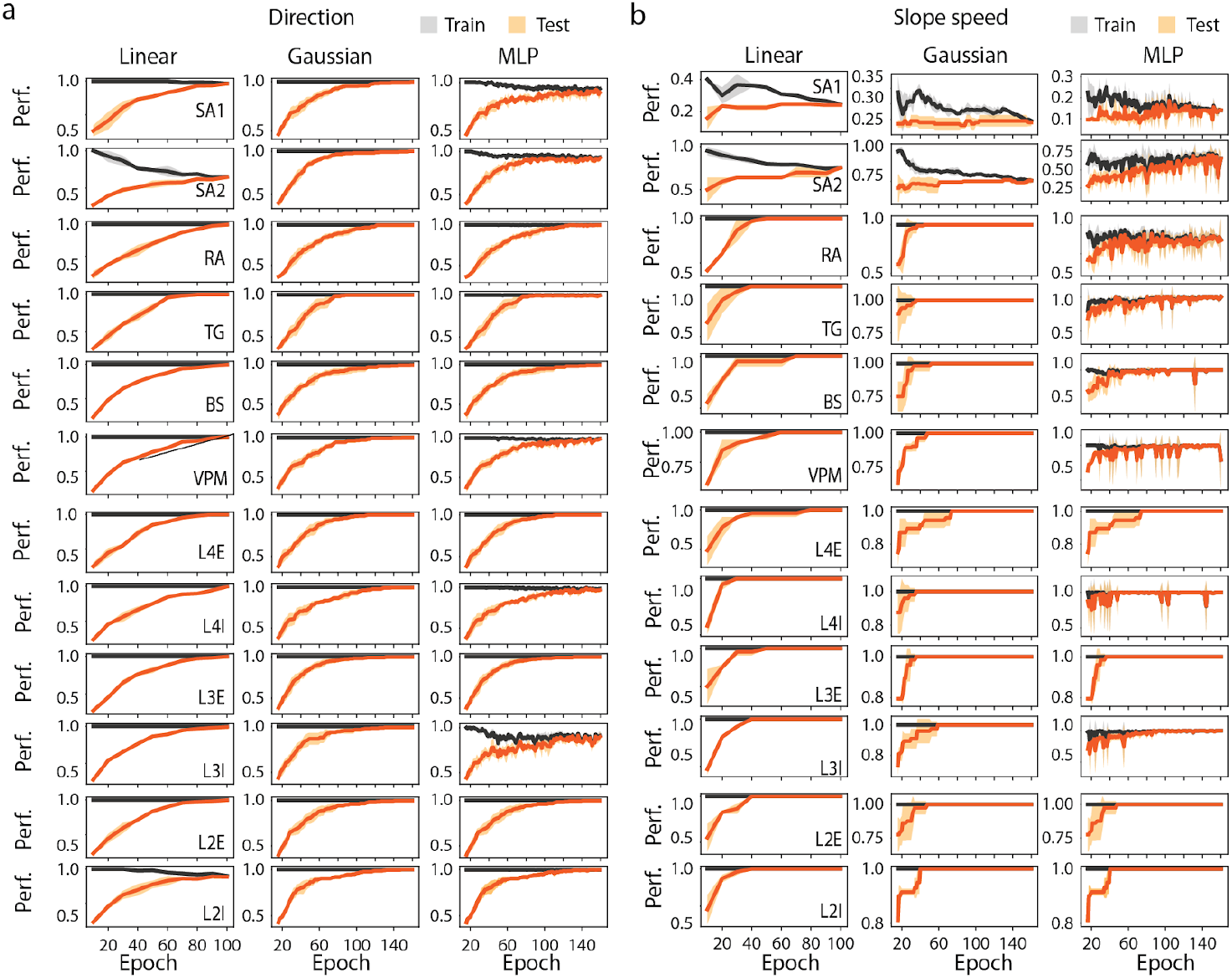
Learning curve for Linear decoder, MLP, and Bayesian when trained to retrieve stimulus **a)** direction and **b)** speed, on whole population activity.

**Supplemental Figure 7.**
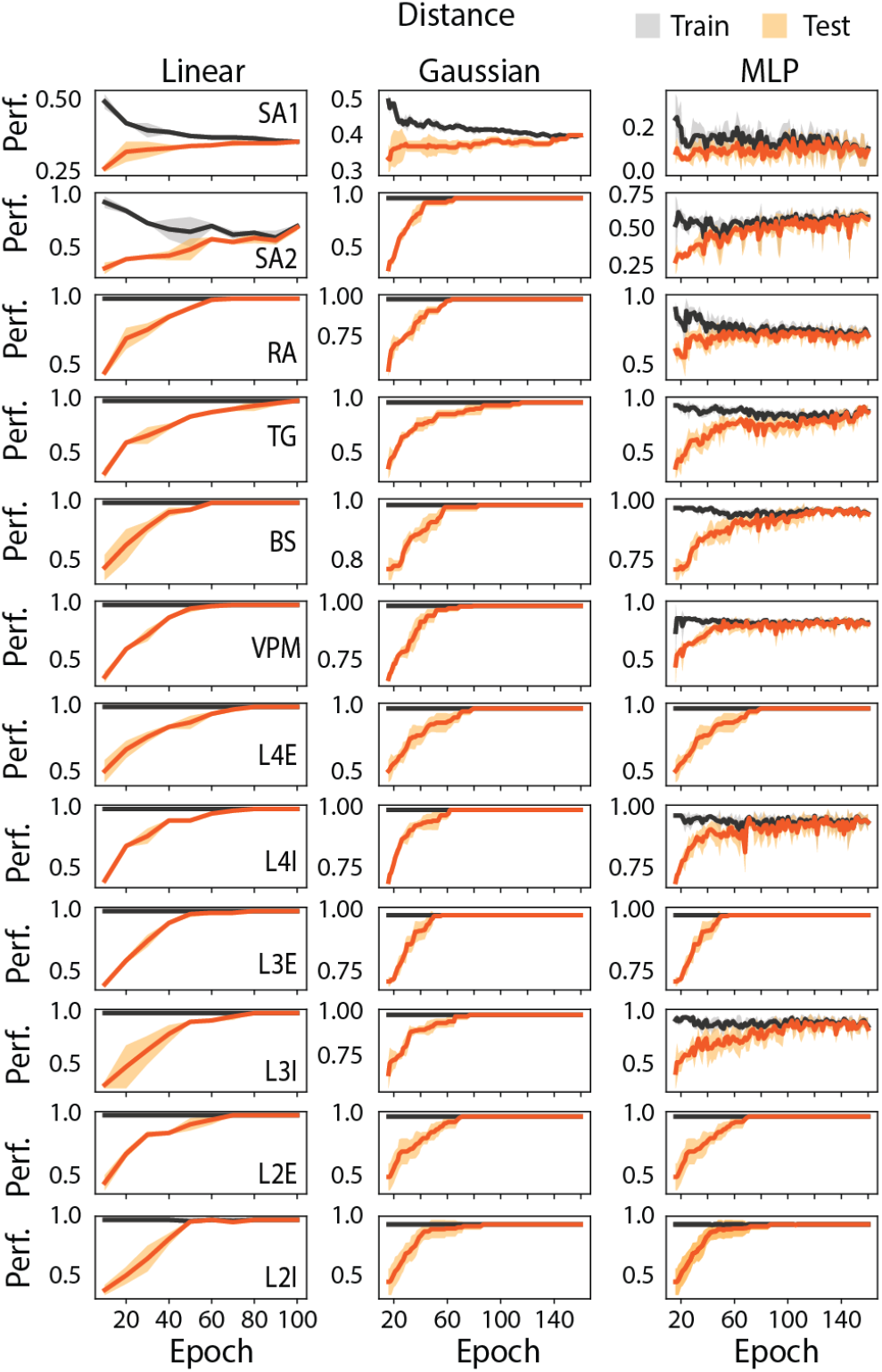
Learning curve for Linear decoder, MLP, and Bayesian when trained to retrieve stimulus distance to whisker basis, on whole population activity.

**Supplemental Figure 8.**
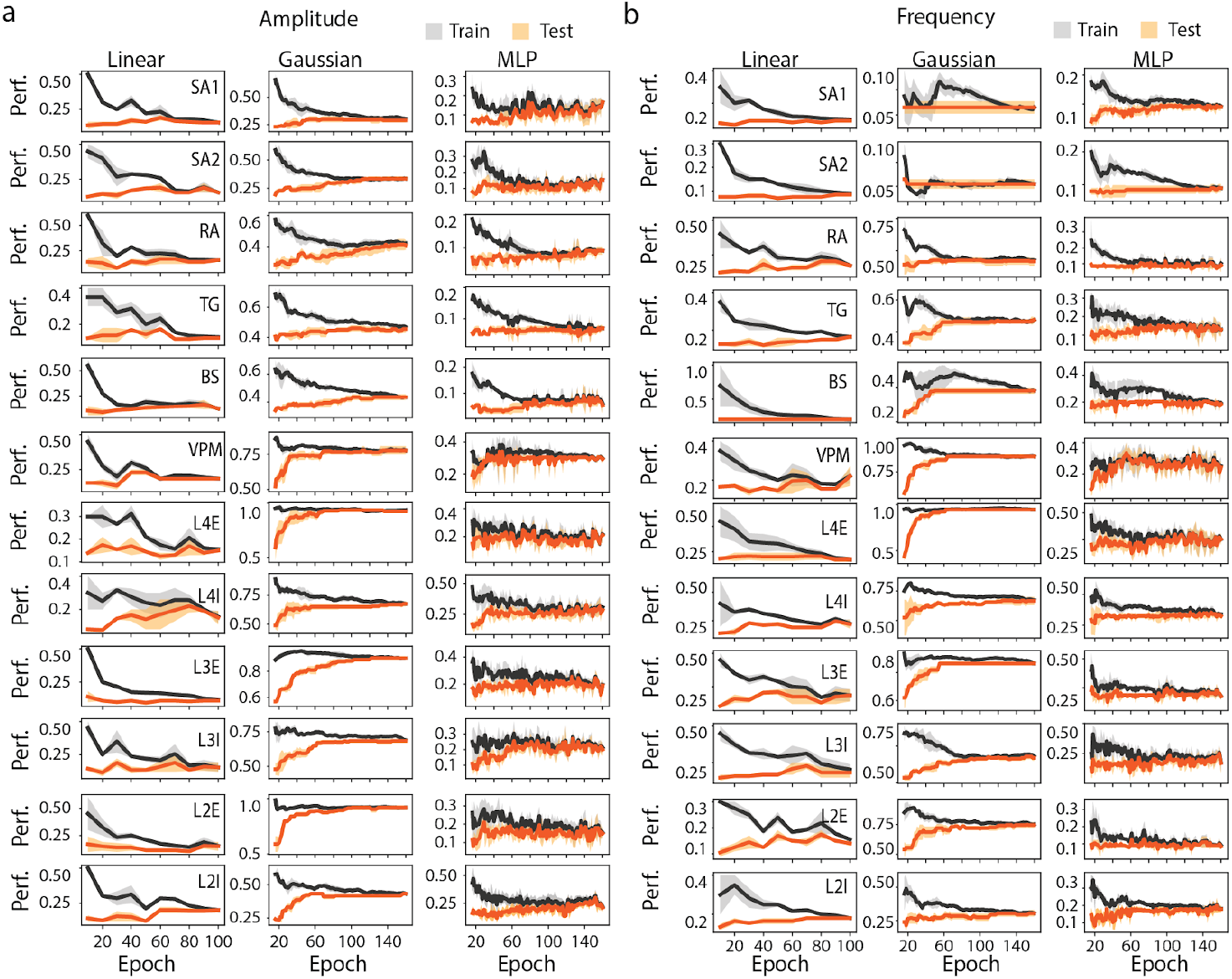
Learning curve for Linear decoder, MLP, and Bayesian when trained to retrieve stimulus **a)** amplitude and **b)** frequency, on the firing rate of the most informative neuron of each population.

**Supplemental Figure 9.**
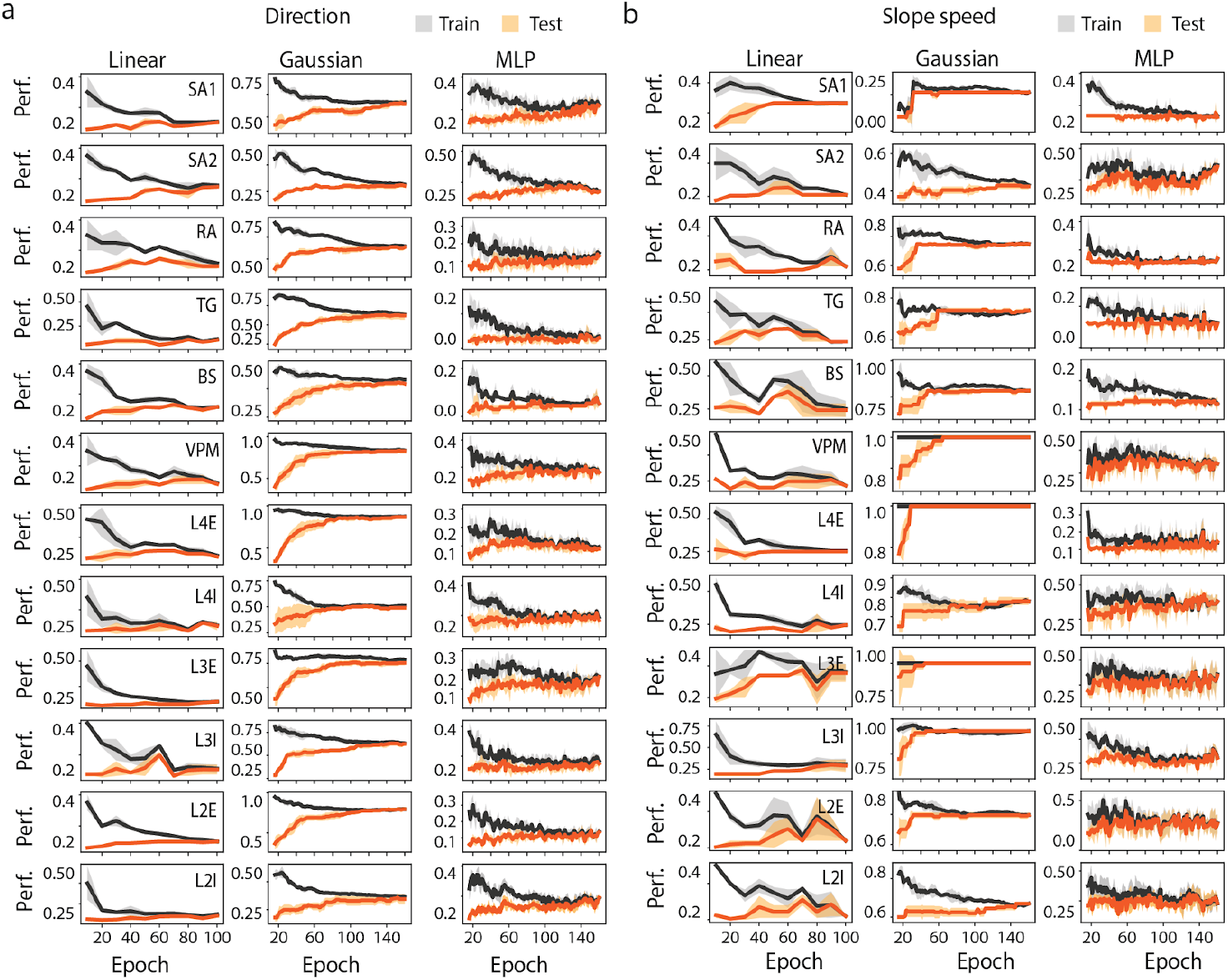
Learning curve for Linear decoder, MLP, and Bayesian when trained to retrieve stimulus **a)** direction and **b)** speed, on the firing rate of the most informative neuron of each population.

**Supplemental Figure 10.**
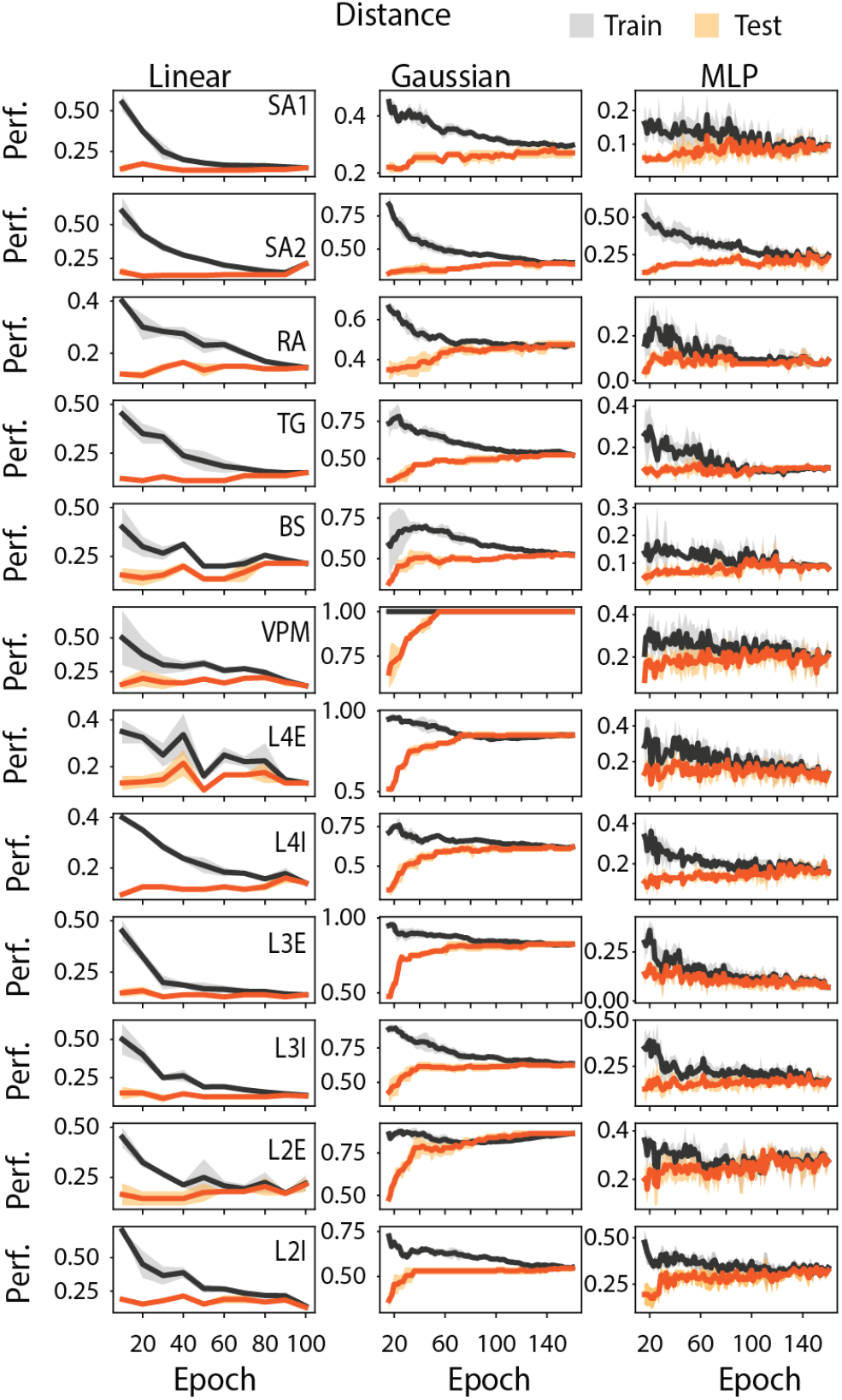
Learning curve for Linear, MLP, and Bayesian decoder when trained to retrieve the stimulus distance to whisker basis, on the firing rate of the most informative neuron of each population.

**Supplemental Figure 11.**
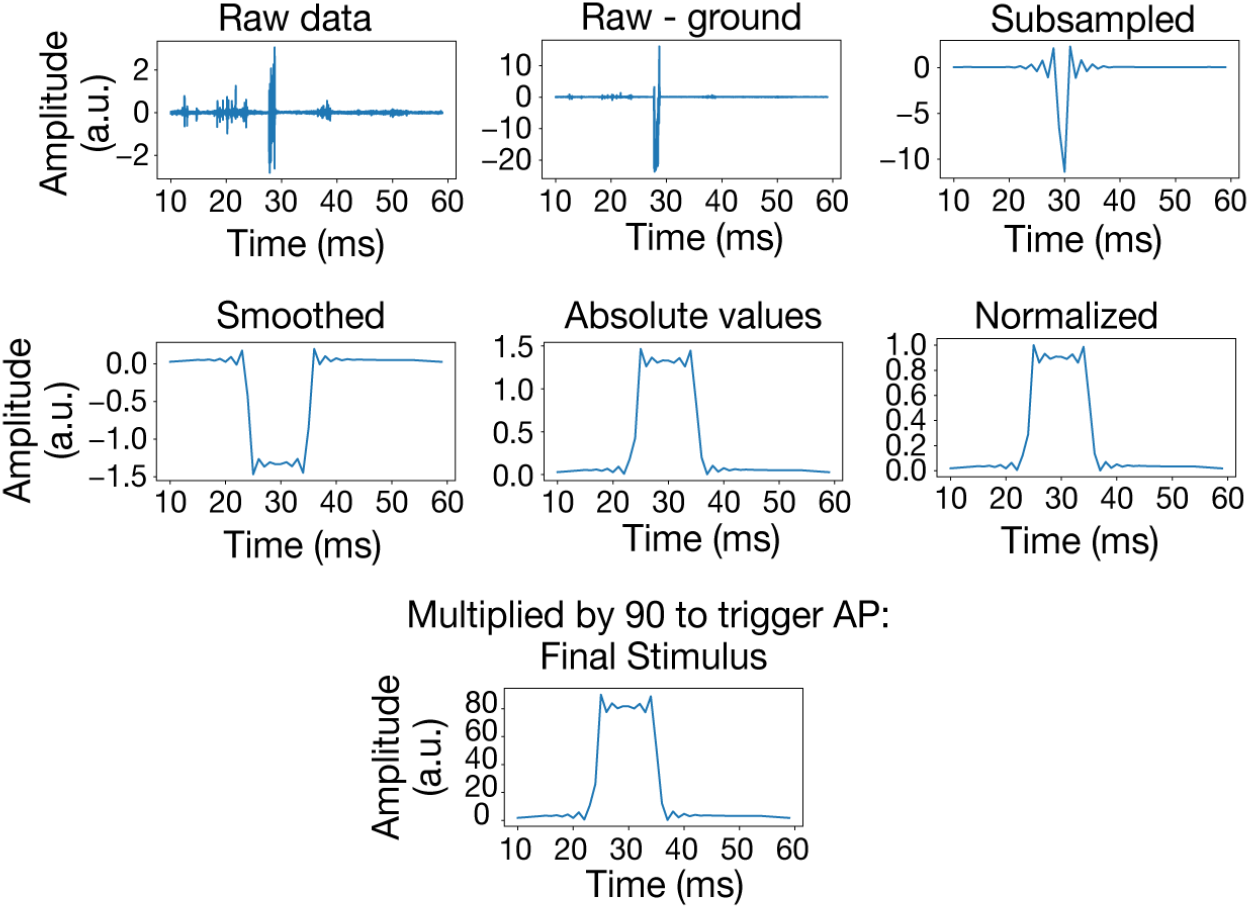
Pipeline, starting from the top row (left) and ending in the bottom row, to process fingertip data prior to network injection. Top row: the resulting stimulus is obtained by substracting the ground data fom the raw, then subsampled to 1kHz. Middle row: It is then smoothed with an average window (11ms), the absolute value is taken, normalized and (bottom row) multiplied by 90 to elicit responses in the model.

**Supplemental Figure 12.**
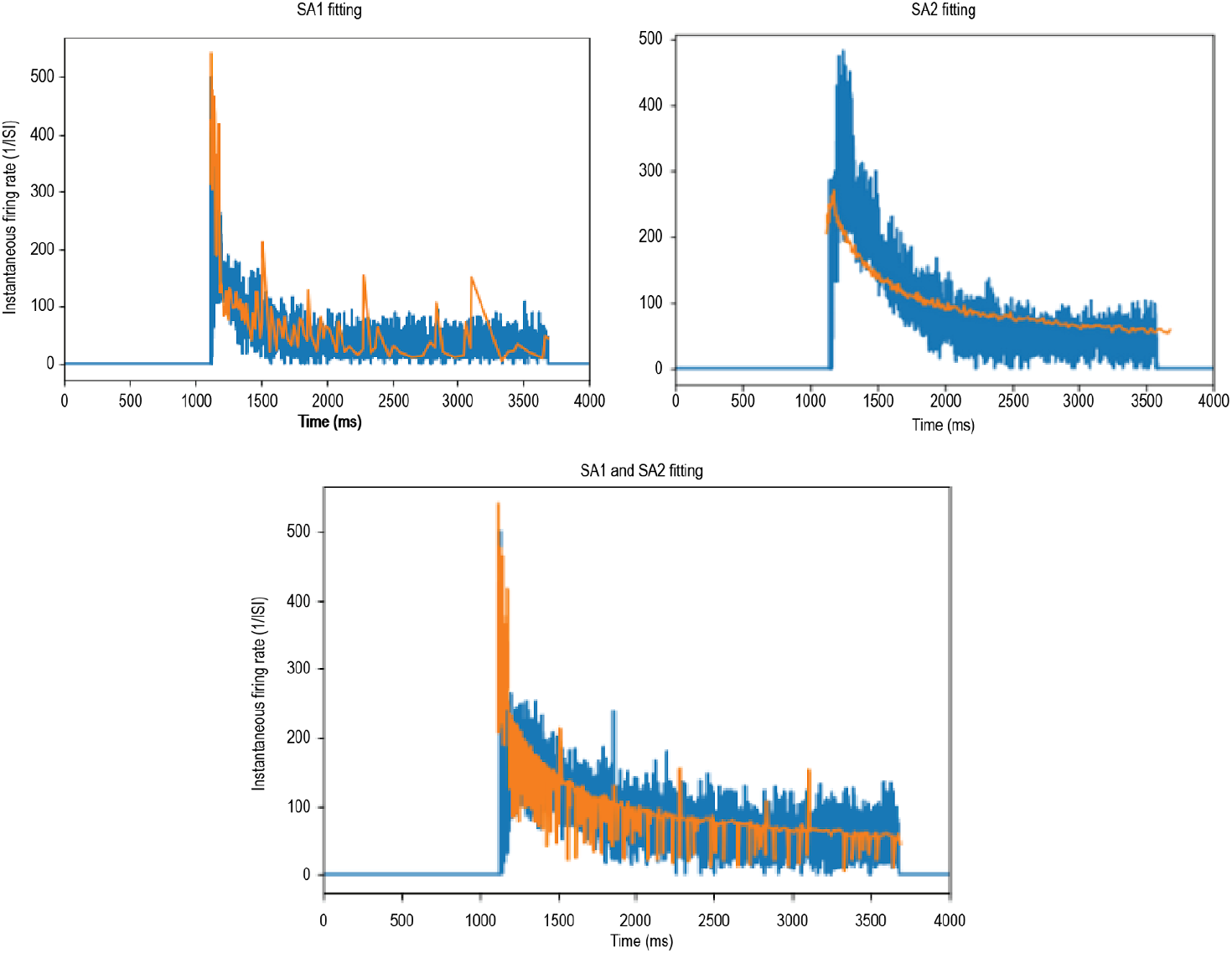
Training results of fitting SA1 and SA2 models. Orange is the model response, blue is the data [19]. A ramp-and-hold stimulus was applied to a whisker. Mechanoreceptor’s activity was recorded from the follicle. The horizontal represents the time in ms and the vertical represents the instantaneous firing rate (1/ISI).

**Supplemental Figure 13.**
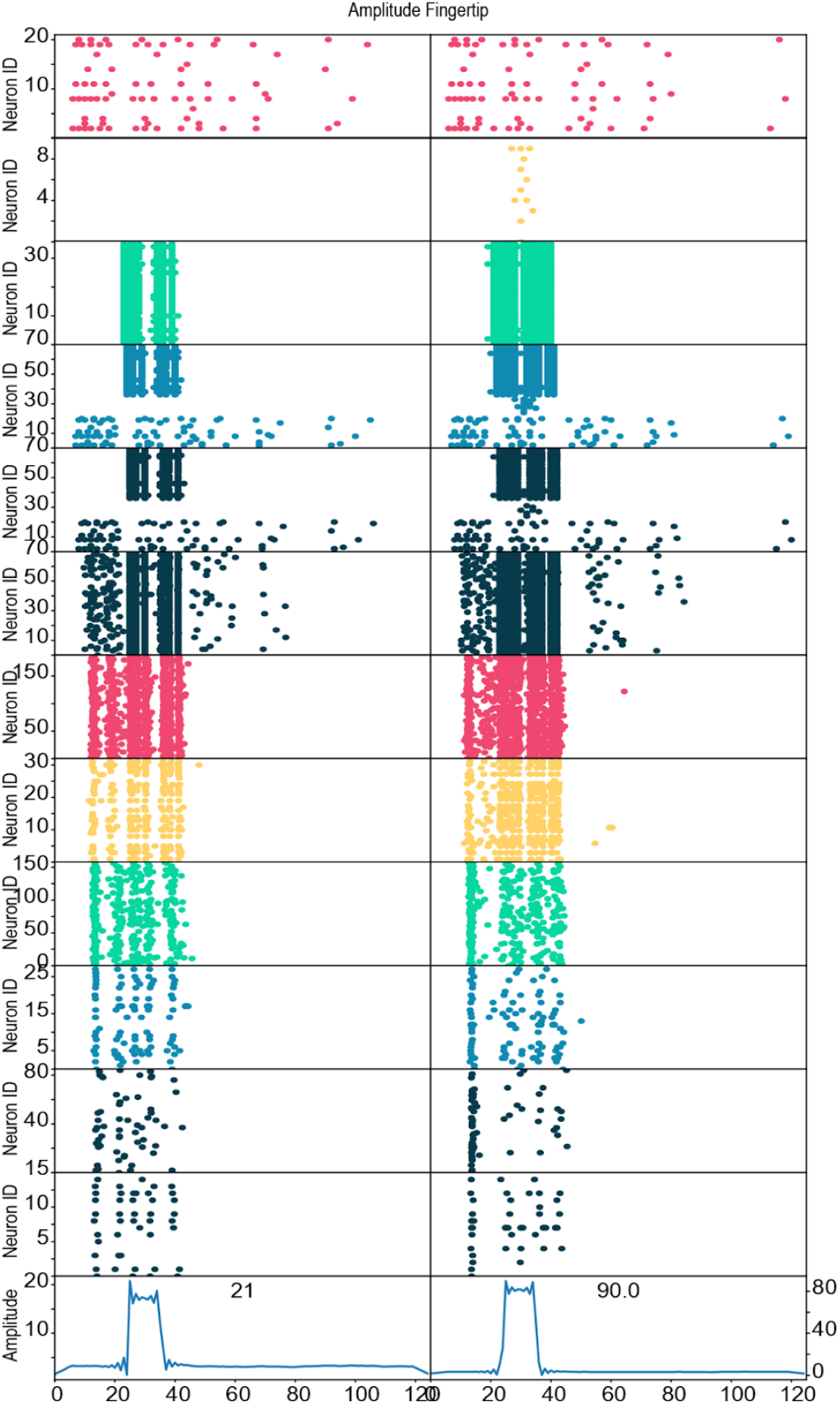
Raster plot across all layers from the model upon application of fingertip displacement input.

**Supplemental Figure 14.**
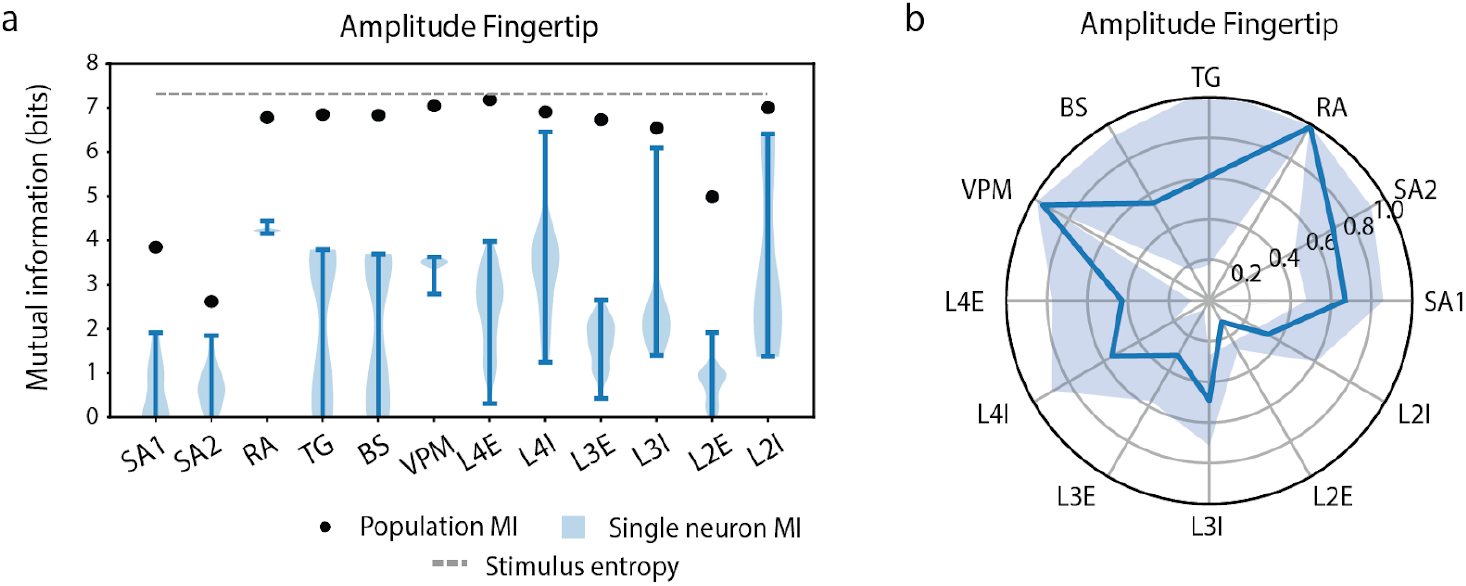
**(a)** Mutual information between single neurons and fingertip stimulus (violin plot) and population and stimulus (blue dot). The blue line corresponds to the entropy of the stimulus. The X-axis is the different populations, Y-axis is the information in bits. **(b)** Pearson correlation coefficient across neurons within a population.

**Supplemental Figure 15.**
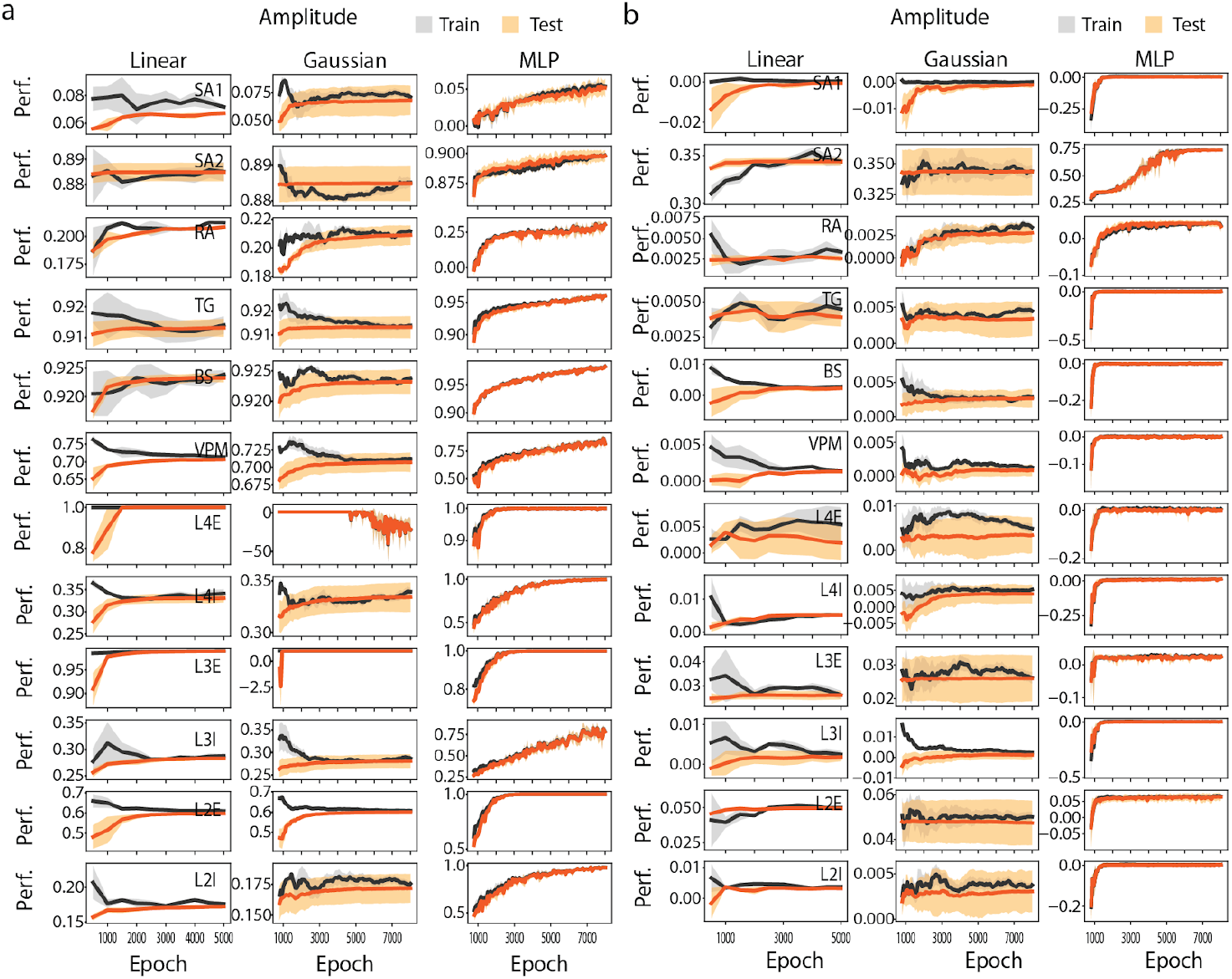
Learning curve for Linear, MLP, and Bayesian decoder when trained to retrieve the fingertip displacement amplitude based on **a)** population activity, and **b)** the activity of the most informative neuron of each population.

